# Essential role of CFAP77-CCDC105-TEX43 subcomplex in connecting axonemal A and B tubules for mammalian sperm motility

**DOI:** 10.1101/2025.02.23.639767

**Authors:** Lan Xia, Guo-Liang Yin, Yu Long, Fei Sun, Bin-Bin Wang, Yun Zhu, Su-Ren Chen

## Abstract

The assembly and physiological function of cilia and flagella depend on the stable association of A and B tubules, which form axonemal microtubule doublets (DMTs). *CFAP77* encodes a core outer junction (OJ) protein within DMTs that is conserved across species and cell types. However, whether and how CFAP77 mediates the connection of B tubules to A tubules of DMTs in mammalian cilia/flagella are unknown. In this study, *Cfap77*-KO mice were generated to reveal that CFAP77 is essential for sperm progressive motility and male fertility. Loss of CFAP77 led to opened B tubules specifically at the OJ regions of axonemal DMTs as revealed by conventional transmission electron microscopy. Cryo-electron tomography further resolved the *in situ* structure of sperm axonemal DMTs from *Cfap77*-KO mice, which exhibited a loss of large filamentous density corresponding to the CFAP77-CCDC105-TEX43 ternary subcomplex at the OJ regions. Additionally, sperm proteomic analysis confirmed that CFAP77 knockout led to the complete loss of this ternary complex. Our work not only explores the physiological role of OJ protein CFAP77 on axonemal A-B tubule connection in mammals but also combines *in situ* structural biology and knockout mice to reveal the underlying structural/molecular mechanism.

## Introduction

Cilia protrude from the surface of almost all eukaryotic cells and have many diverse roles in sensory, motility, and human physiology. The core structure of motile cilia, the axoneme, consists of a pair of microtubule singlets termed as central pair complex (CPC) and nine peripherally positioned doublet microtubules (DMTs), forming a ‘9+2’ architecture^1,2^. Sperm is a type of cell with specialized function and morphology, and their motility is critical for the fertilization^3^. The ‘9+2’ structure is shared by motile cilia and sperm flagella with some differences between species and tissues.

Each DMT has a distinctive structure with a complete ring of 13 protofilaments (the A tubule) and an incomplete ring of 10 protofilaments (the B tubule)^1^. The heads of dynein arms as well as the base of radial spoke and nexin-dynein regulatory (N-DRC) complex attach to the A tubules while the tails of dynein and N-DRC connect the neighboring B tubules^2^. Connections of A and B tubules is expected to be critical for axonemal stability and cilia/flagella motility. A long-standing question is how B tubules tightly attach to A tubules within DMTs.

Inner junction (IJ) at A01-B10 and outer junction (OJ) proteins at A11-A12 and B01-B02 are suggested to mediate this connection^1,2^. Recently, we and others explore the high-resolution structure of human and mouse sperm flagella, revealing many core and sperm-specific microtubule inner proteins (MIPs)^6-9^. CFAP20 and PACRG are identified as core components of the IJ while CFAP77 forms the OJ connection between A and B tubules^5,10-11^. Their expression are conserved in cilia/flagella across many species including *Chlamydomonas*, mouse, and humans^6,9^. In zebrafish and *C. elegans*, CFAP20 maintains the structural integrity of IJ and is required for cilia motility^12^. Knockout of CFAP77 moderately reduces *Tetrahymena* cell motility and partially destabilizes the OJ^5^. However, their effects on mammalian cilia/flagella function and the underlying structural/molecular mechanism of IJ/OJ stability are largely unclear.

Structural biology technologies, such as cryo-electron microscopy (cryo-EM) and *in situ* cryo-electron tomography (cryo-ET) bring our understating of the axoneme to the molecular level. By a combination with knockout animal models, these structural biology technologies will further provide direct visual evidence to explore the structural/molecular mechanisms. For example, deficiency of CFAP47 leads to sperm motility problem and human/mouse infertility without understanding the molecular mechanism^13^. By resolving the *in situ* high-resolution structures of sperm axoneme, we found that CFAP47 is specifically localized at the bridge of sperm CPC^14^. Missing density (e.g., CFAP47 and SPAG6) in the bridge region is clearly revealed by resolving *in situ* structure of sperm CPC from *Cfap47*-KO mice, representing as the structural/molecular basis of sperm motility abnormility^14^.

In this study, we generated a *Cfap77*-KO mouse strain and found that knockout mice were male infertile with defects in sperm motility. Intriguingly, loss of CFAP77 led to a disconnection of A-B tubules specifically at the OJ regions by transmission electron microscopy analysis of axonemal ultrastructures. Cryo-ET further resolved the *in situ* structure of the mutant axonemal DMTs, which exhibited a loss of large filamentous density corresponding to the CFAP77-CCDC105-TEX43 ternary subcomplex at the OJ regions. Absence of this ternary complex represented the initial event underlying disconnection of A and B tubules at the OJ sites, axoneme instability, impaired sperm progressive motility, and male infertility in *Cfap77*-KO mice.

## Results

### CFAP77-CCDC105-TEX43 subcomplex at the OJ regions of sperm DMTs

Recently, we and others resolved the near-atomic-level structure of DMTs from mouse sperm^6-9^, which provides a valuable source to further explore the function of individual structural component of DMTs. CFAP77, CCDC105, and TEX43 caught our attention because they are localized at the OJ regions in both isolated sample (PDB entry 8IYJ) and *in situ* sample (PDB entry 8I7R), which may play important roles in the connection of B tubules to A tubules within DMTs (Fig. 1A). In mouse sperm, CFAP77 binds to A11-A12 and B01-B02 protofilaments in a 16 nm repeat. CCDC105 resides between the protofilaments A11 and A12, assembling into a long filament by interacting with the neighbor CCDC105 molecules. It seems that CCDC105 filaments contact individual CFAP77 protein into a stable structure to connect A and B tubules. TEX43 is located at the junction site of two CCDC105 proteins, implying a role to reinforce of CCDC105 filaments (Fig. 1A and B).

**Fig 1.**
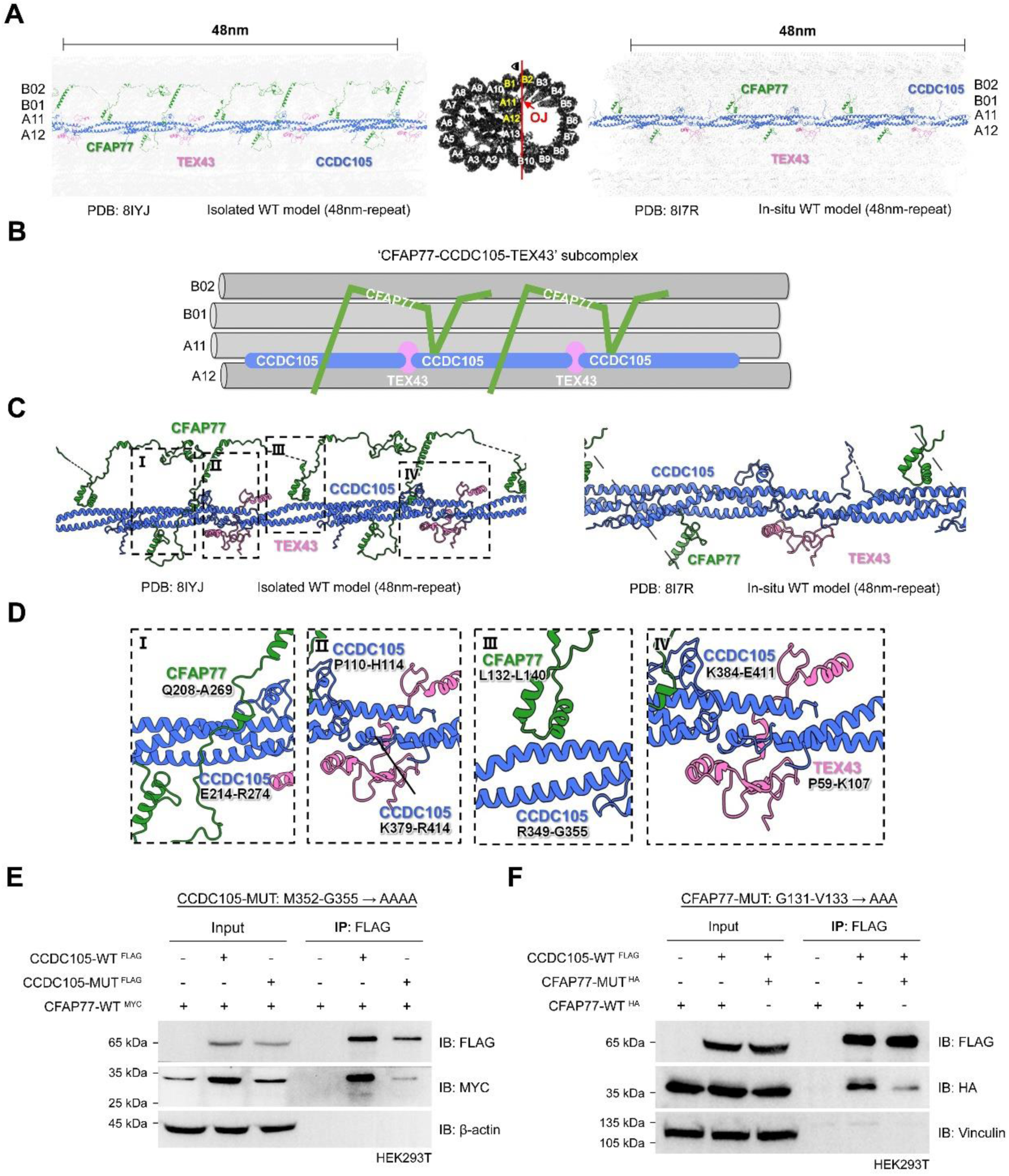
CFAP77-CCDC105-TEX43 subcomplex at the OJ sites of sperm axoneme. **(A)** Structural models of CFAP77-CCDC105-TEX43 subcomplex at the OJ regions of mouse sperm axoneme were shown with a 48 nm length of vertical section in the isolated DMT (PDB: 8IYJ) and in-situ DMT (PDB: 8I7R). CFAP77 was resided between A11-A12 and B01-B02. OJ, outer junction. **(B)** Simplified schematic diagram to illustrate the connection of individual CFAP77 by CCDC105 bundles and organization of CCDC105 bundles by TEX43. CCDC105 and TEX43 were localized at A11-A12. **(C)** The interaction interfaces of CFAP77-CCDC105-TEX43 subcomplex were shown in isolated mouse sperm DMT model (PDB: 8IYJ) and in-situ mouse sperm DMT model (PDB: 8ITR). **(D)** The interaction interfaces of CFAP77-CCDC105 (I), CCDC105-CCDC105 (II), CCDC105-CFAP77 (III) and CCDC105-TEX43 (IV) which were indicated in **(C)**. Residues at the interaction interfaces were labeled. **(E)** The co-immunoprecipitation of MYC-tagged CFAP77 by FLAG-tagged WT-CCDC105 and mutant (MUT)-CCDC105 in HEK293T cells. For CCDC105 mutant plasmid, 352-355 amino acids were mutated into AAAA. β-actin served as the internal control. **(F)** The interacting ability of FLAG-tagged CCDC105 with HA-tagged WT-CFAP77 or MUT-CFAP77 in HEK293T cells. For CFAP77 mutant plasmid, 131-133 amino acids were mutated into AAA. Vinculin served as the internal control.

There are rich and close interactions among CFAP77, CCDC105 and TEX43^7,9^, making them resemble a relatively independent subcomplex within DMTs (Fig. 1C). CCDC105 features two helical bundles: one bundle is wrapped by the glutarnine (Q)208-alanine (A)269 region of CFAP77, while the other binds to the leucine (L)132-L140 region of CFAP77 (Fig. 1D). Additionally, the proline (P)59-lysine (K)107 region of TEX43 interacts with the interface of two adjacent CCDC105 proteins (Fig. 1D). To validate the interactions in this ternary subcomplex, we mutated the key residues at one of the interaction interfaces between CFAP77 and CCDC105. Methionine (M)352-glycine (G)355 residues of CCDC105 were mutated into alanine (A)AAA and G131-valine (V)133 residues of CFAP77 were mutated in to AAA, respectively. By co-immunoprecipitation (Co-IP) analysis, we showed that the interaction between CFAP77 and CCDC105 was obviously attenuated after mutations of those critical contact sites (Fig. 1E and F). Together, these results pose a hypothesis that CFAP77-CCDC105-TEX43 subcomplex at the OJ regions is critical for DMTs A-B tubule connection and sperm motility.

### *Cfap77*-KO mice are male infertile and produce sperm with motility defects

Knockout of CFAP77 moderately reduces *Tetrahymena* cell motility and partially destabilizes the OJ^5^. To reveal the physiological role of CFAP77 in mammals, we generated a first *Cfap77*-KO mouse strain by applying CRISPR/Cas9 technology. Exons 2 and 3 of transcript *Cfap77*-202 (ENSMUST00000157048) was selected as the knockout region to generate a 30,988 bp deletion (Fig. 2A). By Western blot analysis, CFAP77 expression was confirmed to be completely absent in the sperm samples of knockout mice (Fig. 2B). *Cfap77*-KO mice were healthy and developed normally; however, the fertility test showed that *Cfap77*-KO male mice were infertile (Fig. 2C). In vitro fertilization (IVF) assay also indicated that the rate of 2-cell embryos was significantly lower in the groups using the sperm from *Cfap77-*KO mice than those from wild-type (WT) mice (Fig. 2D and E). Histological examination of testis and cauda epididymis from *Cfap77-*KO mice further revealed a normal process of spermatogenesis and sperm production (Fig. S1). To explore the reason of infertility of *Cfap77-*KO mice, we next examined the semen characteristics. As revealed by Papanicolaou stain, the morphology of sperm from *Cfap77*-KO mice was found to be generally normal (Fig. 2F). Sperm count was also similar between *Cfap77*-KO mice and WT mice (Fig. 2G), whereas progressive sperm motility was significantly attenuated in *Cfap77*-KO mice (Fig. 2H). By sperm motion tracking and statistical analysis, we found that the percentage of sperm showing S-shape, O-shape, and shake/silent movement was significantly increased in *Cfap77*-KO mice as compared with WT mice (Fig. 2I and Movies 1-2). Collectively, the animal studies indicate that CFAP77 is critical for progressive sperm motility and male fertility in mammals.

**Fig 2.**
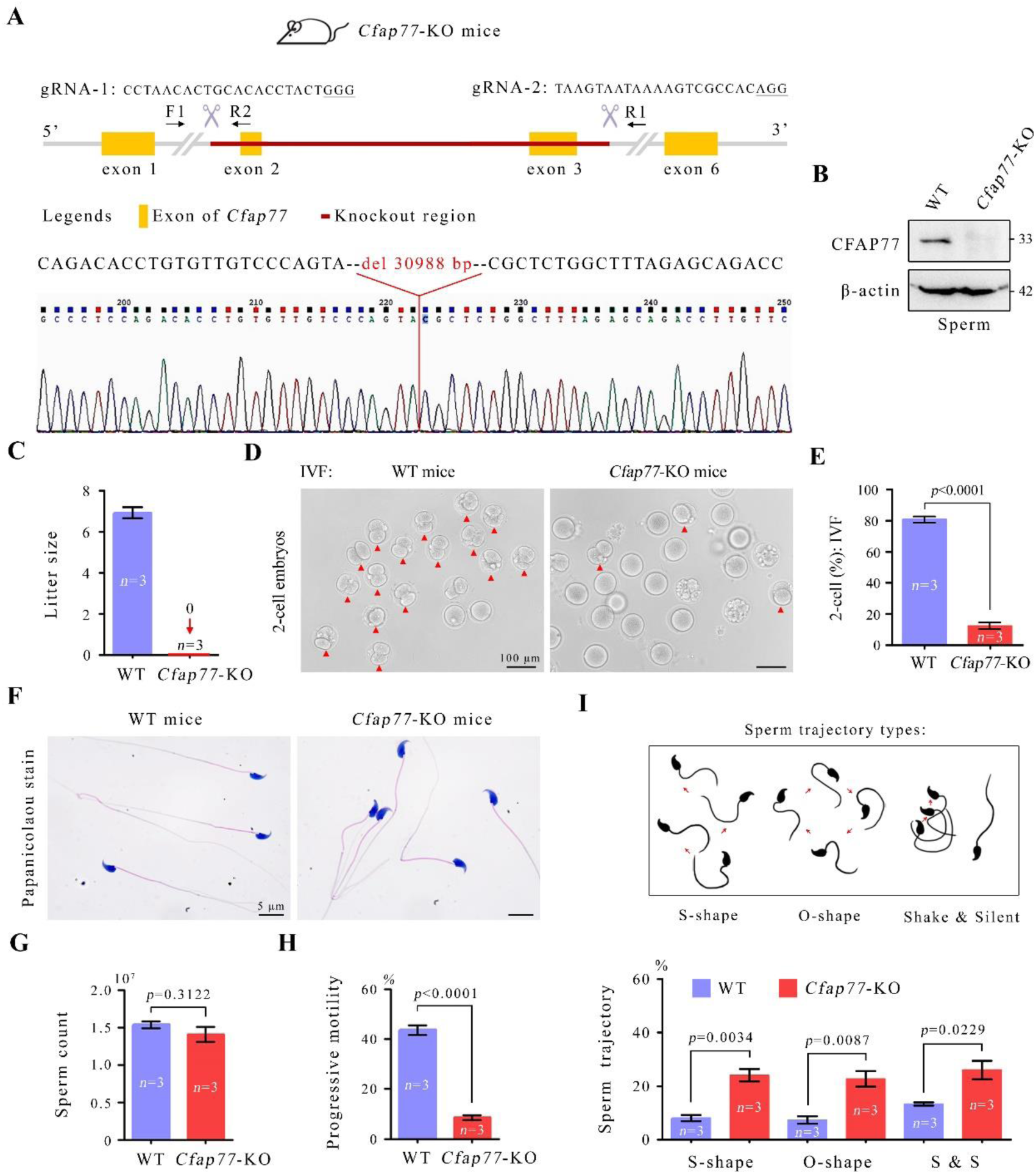
*Cfap77*-KO mice were sterile due to defective sperm motility. **(A)** Genomic features and knockout strategy of mouse *Cfap77* by CRISPR/Cas9 technology. A 30,988 bp deletion (targeting exons 2-3) in the *Cfap77* gene was confirmed by Sanger sequencing. **(B)** Western blot analysis of CFAP77 protein expression in sperm samples from WT and *Cfap77*-KO mice. β-actin served as an internal control. **(C)** The litter size of WT female mice was calculated between WT male groups and *Cfap77*-KO male groups. Error bars represent SEM (*n*=3). Statistical analysis could not be performed due to all data in *Cfap77*-KO male groups are 0. **(D)** Representative 2-cell embryos from in vitro fertilization (IVF) experiments. Red arrowheads indicated the 2-cell embryos. Scale bars, 100 μm. **(E)** The percentage of 2-cell embryos in the group using sperm from *Cfap77*-KO mice or using sperm from WT mice. Student’s *t* test; error bars represent SEM (*n*=3). **(F)** Papanicolaou staining of sperm from WT mice or *Cfap77*-KO mice. Scale bars, 5 μm. **(G)** Sperm count of WT mice and *Cfap77*-KO mice. Student’s *t* test; error bars represent SEM (*n*=3). **(H)** Progressive motility of sperm from WT mice and *Cfap77*-KO mice. Student’s *t* test; error bars represent SEM (*n*=3). **(I)** Schematic diagram illustrating three major types of sperm trajectory. The percentage of sperm showing S-shape, O-shape, and shake & silent types of motility between WT mice and *Cfap77*-KO mice. Student’s *t* test; error bars represent SEM (*n*=3).

### Open DMT-B tubules at the OJ regions in sperm of *Cfap77*-KO mice

To investigate the structural basis for impaired progressive sperm motility in *Cfap77*-KO mice, we examined the ultrastructure of sperm flagellar axoneme by traditional transmission electron microscopy (TEM). The ultrastructures of acrosome and other sperm regions were normal in *Cfap77*-KO mice (Fig. S2A). Accessory structures of axonemes such as mitochondrial sheath in the mid-piece and fibrous sheath in the principal piece of sperm flagella were also indistinguishable between *Cfap77*-KO mice and WT mice (Fig. S2A). Focusing on axoneme, there was no discernible deficiency of the CPC, radial spokes, dynein arms, and N-DRC in *Cfap77*-KO mice. The disconnection of A and B tubules within DMTs represented as the only ultrastructural defect after the loss of CFAP77 (Fig. 3). Sperm from WT mice exhibited well-organized “9+2” axonemes and DMT-B tubules were closely attached to the DMT-A tubules even after sperm capacitation for 1 hour (Fig. 3A). By contrast, a large proportion of sperm in *Cfap77*-KO mice showed defects specifically at the A and B tubule connection site (open DMT-B tubule at the OJ regions) (Fig. 3B). In axonemes of *Cfap77*-KO mice, the disconnection of A and B tubules was mainly distributed to DMTs 1, 5, 6, 9 and 4 (Fig. S2B). By statistical analysis, the percentage of axonemes with open DMT-B tubules was found to be significantly increased after the loss of CFAP77 protein (6.67%±1.453% in WT mice and 45.33%±3.480% in *Cfap77*-KO mice) (Fig. 3C). In *Cfap77*-KO mice, this ratio showed no obviously different between mid-piece and principal piece of sperm flagella (Fig. S2C). Intriguingly, the percentage of disconnection of A and B tubules was further increased in *Cfap77*-KO mice after sperm capacitation for 1 hour (72.00%±4.141%), suggesting that the instability of axonemes was increasing during sperm hyperactivation (Fig. 3C). Collectively, the ultrastructural analysis clearly indicates that detachment of A-B tubules within DMTs is the underlying reason of the disruptive sperm progressive motility in *Cfap77*-KO mice. The phenotype of open DMT-B tubules specifically at the OJ regions in *Cfap77*-KO mice is also perfectly consistent with the expression evidence that CFAP77 is selectively localized at the OJ of DMTs.

**Fig 3.**
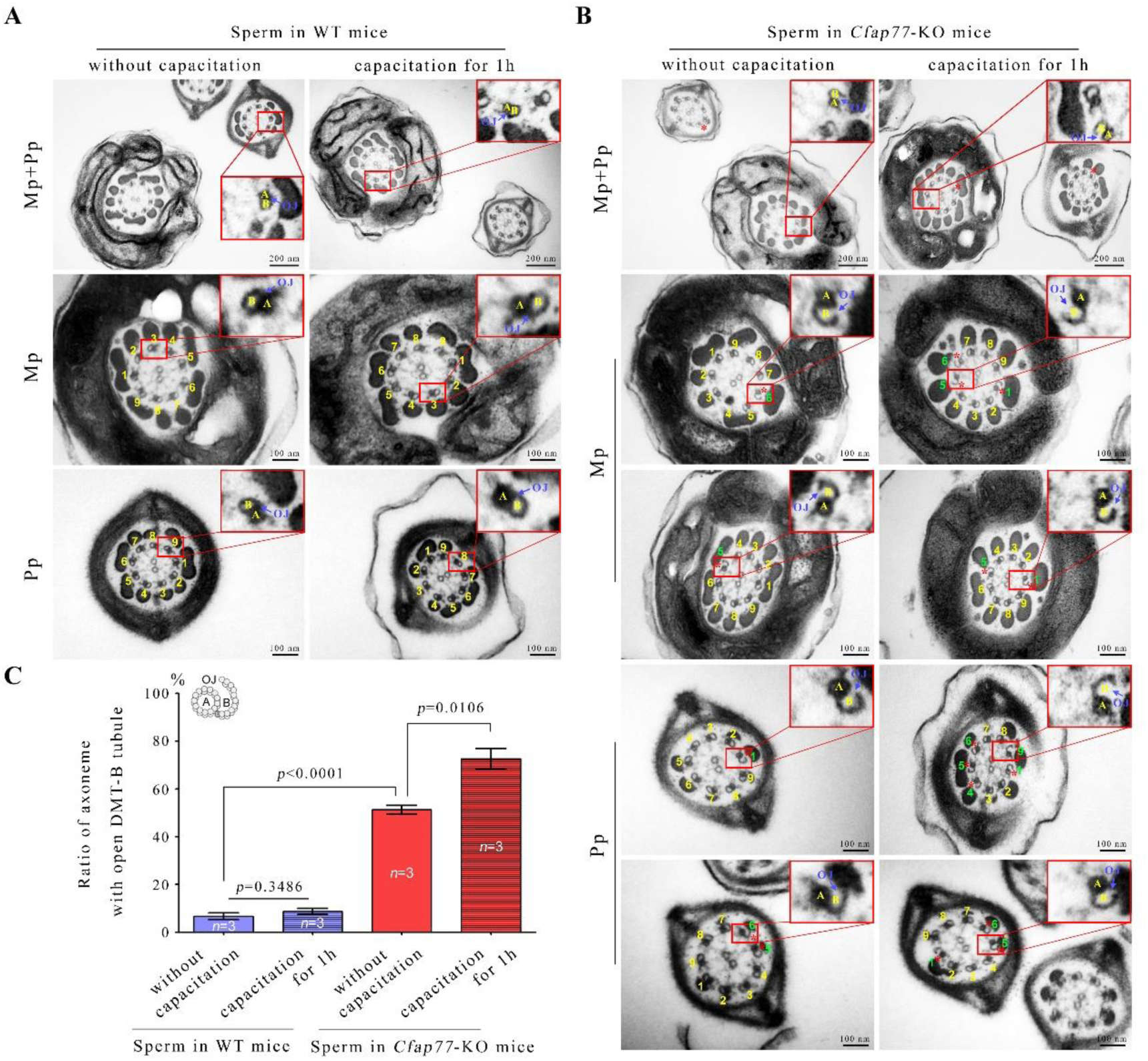
Open DMT-B tubules of sperm axonemes in *Cfap77*-KO mice. **(A)** Transmission electron microscope (TEM) analysis of sperm axoneme (9+2) from WT mice. Sperm collected from cauda epididymis were directly fixed or underwent capacitation in TYH medium for 1 hour and then fixed. Each DMT was numbered. Scale bars, 200 nm or 100 nm. **(B)** TEM images of sperm axonemes from *Cfap77*-KO mice. Principal piece (Pp) and mid-piece (Mp) of sperm flagella were shown and a single DMT was zoomed in. Each DMT was numbered. Green numbers and red stars indicated the OJ regions where open DMT-B tubules exist. Scale bars, 200 nm or 100 nm. **(C)** The percentage of sperm axonemes with open DMT-B tubules was calculated between WT mice and *Cfap77*-KO mice, as well as uncapacitation and capacitation subgroups. Student’s *t* test; error bars represent SEM (*n*=3).

### In-cell structural insight of sperm DMTs in *Cfap77*-KO mice

To investigate the impact of CFAP77 deletion on the structural assembly of sperm DMTs at a higher resolution, we frozen the sperm from *Cfap77*-KO mice on the grid and milled the sample to approximately 200 nm thick using cryo-focused ion beam thinning (cryo-FIB) (Fig. S3). We collected cryo-ET data targeting the sperm tail, and resolved the *in situ* structure of DMTs using sub-tomogram averaging (STA) approach (Fig. S4). During particle picking and data processing, we found that DMT particles showed notable structural heterogeneity, indicating that the ultrastructure of sperm DMTs in *Cfap77*-KO mice was compromised. This is consistent with our conventional TEM observations that approximately 45% of DMT-B tubules was opened at the OJ regions in sperm from *Cfap77*-KO mice (Fig. 3). After discarding many particles through 3D classification, we obtained the 8 nm repeat structure of DMTs with a resolution of 24 Å (Fig. S5). This sperm DMTs from *Cfap77*-KO mice showed generally the same overall shape compared to the structure of DMTs in WT mice^9^ (Fig. 4A), except for the absence of a significant density near protofilaments A11 and A12 (Fig. 4B). From the difference map, we could also confirm that the major missing part of the DMTs structure was a long fibrous density extending longitudinally along the tubulin wall of the ribbon (Fig. 4B). Then, we fitted the atomic models of sperm DMTs structure of WT mice^9^ into the density map of sperm DMTs of *Cfap77*-KO mice (Fig. 4C and Fig. S5). Notably, CFAP77, CCDC105, and TEX43 were identified to be the missing densities in the sperm DMT-B tubules of *Cfap77*-KO mice (Fig. 4C). The CFAP77 corresponding density was absent due to the knockout of *Cfap77* gene, as expected, but the loss of CCDC105 and TEX43 highlights the essential role of CFAP77 for the formation of CFAP77-CCDC105-TEX43 ternary complex. Collectively, our in-cell structural study provides direct visual evidence that loss of CFAP77-CCDC105-TEX43 subcomplex at the OJ regions represents the initial event of open DMT-B tubules of sperm axonemes in *Cfap77*-KO mice.

**Fig 4.**
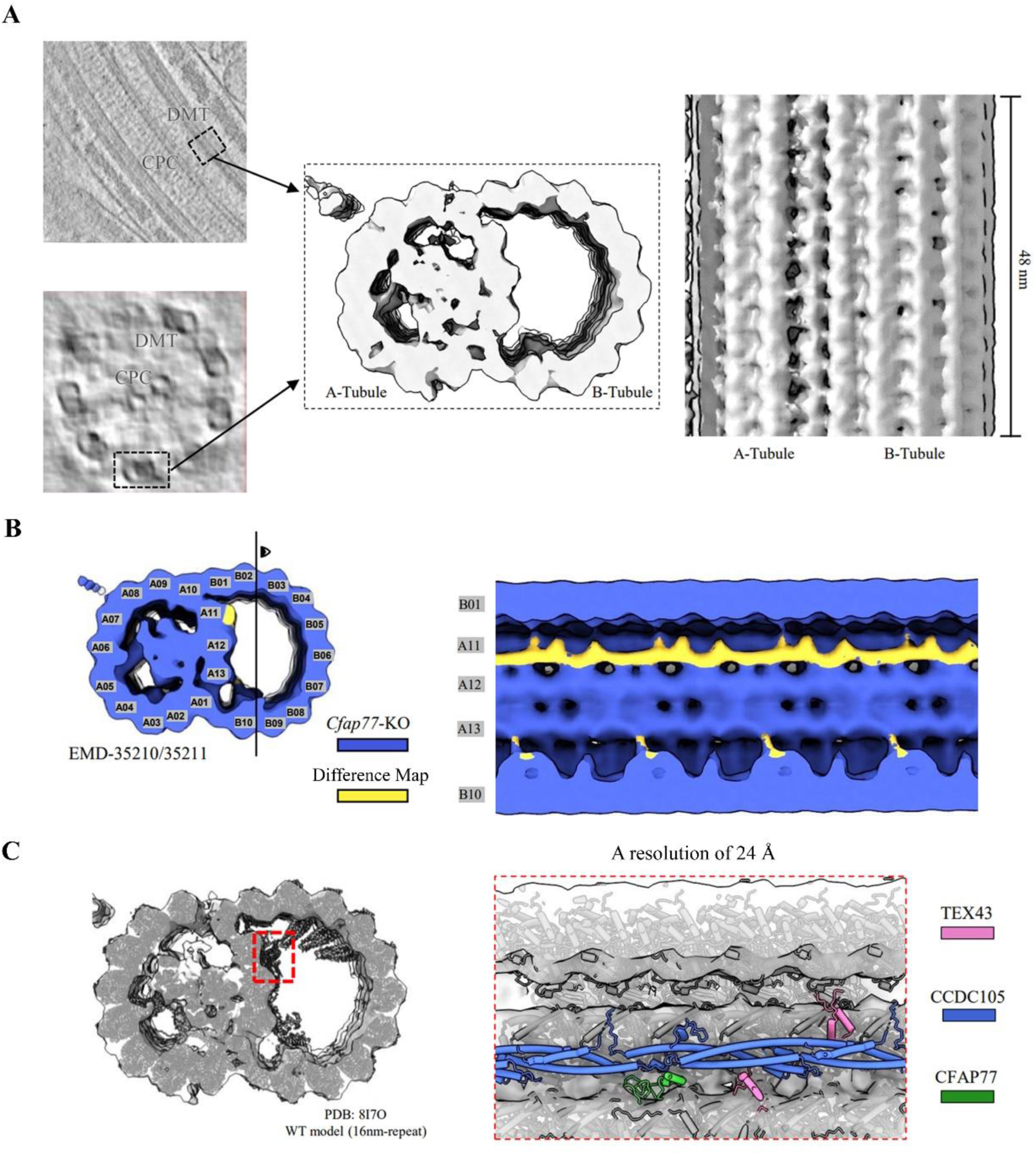
Overall architecture of sperm axonemal DMTs in *Cfap77*-KO mice. **(A)** Schematic representation of in-cell structural determination of sperm axonemal DMTs from *Cfap77*-KO mice. Side view and transverse sectional view of mouse sperm axoneme were shown in the tomogram slices. The cryo-EM map of DMTs with an 8 nm repeat was obtained by sub-tomogram analysis. DMT, microtubule doublets; CPC, central pair complex. **(B)** Loss of density in sperm axonemal DMTs from *Cfap77*-KO mice. The transverse sectional view of DMTs and the side view of B tubule lumen were shown. The lost density (yellow color) was obtained by subtracting the density map of *Cfap77*-KO DMTs from that of the WT DMTs (EMD-35210/35211). **(C)** The atomic model of WT DMTs (PDB: 8I7O) fitted into the density map of *Cfap77*-KO DMTs. The *Cfap77*-KO DMTs showed a loss of density corresponding to CFAP77-CCDC105-TEX43 subcomplex near protofilaments A11 and A12.

### Missing CCDC105 and TEX43 proteins after CFAP77 deletion

Given that the sperm count and morphology are highly similar between *Cfap77*-KO mice and WT mice, mass spectrometry were further applied to quantitatively identify the sperm proteomics of WT and *Cfap77*-KO mice (*n*=3 each group) (Fig. 5A). A cut-point of 5-fold change and a *p*-value (student’s *t* test) less than 0.05 was selected as the screening criteria. Compared with WT group, CCDC105 and TEX43 proteins were identified to be significantly downregulated (more than 20-fold reduction) in sperm samples of *Cfap77*-KO mice. This proteome discovery is perfectly consistent with our *in situ* structural analysis of sperm DMTs from *Cfap77*-KO mice showing missing CFAP77-CCDC105-TEX43 subcomplex at the OJ regions (Fig. 4). By Western blot analysis, we further confirmed that the expression CCDC105 (Fig. 5B) and TEX43 (Fig. 5C) was almost undetectable in both testis and sperm samples from *Cfap77*-KO mice. In contrast, the mRNA levels of *Ccdc105* (Fig. 5D) and *Tex43* (Fig. 5E) were unaltered after the deletion of CFAP77, indicating that CFAP77 affects the expression/stability of CCDC105 and TEX43 at the protein level rather than the mRNA level.

**Fig 5.**
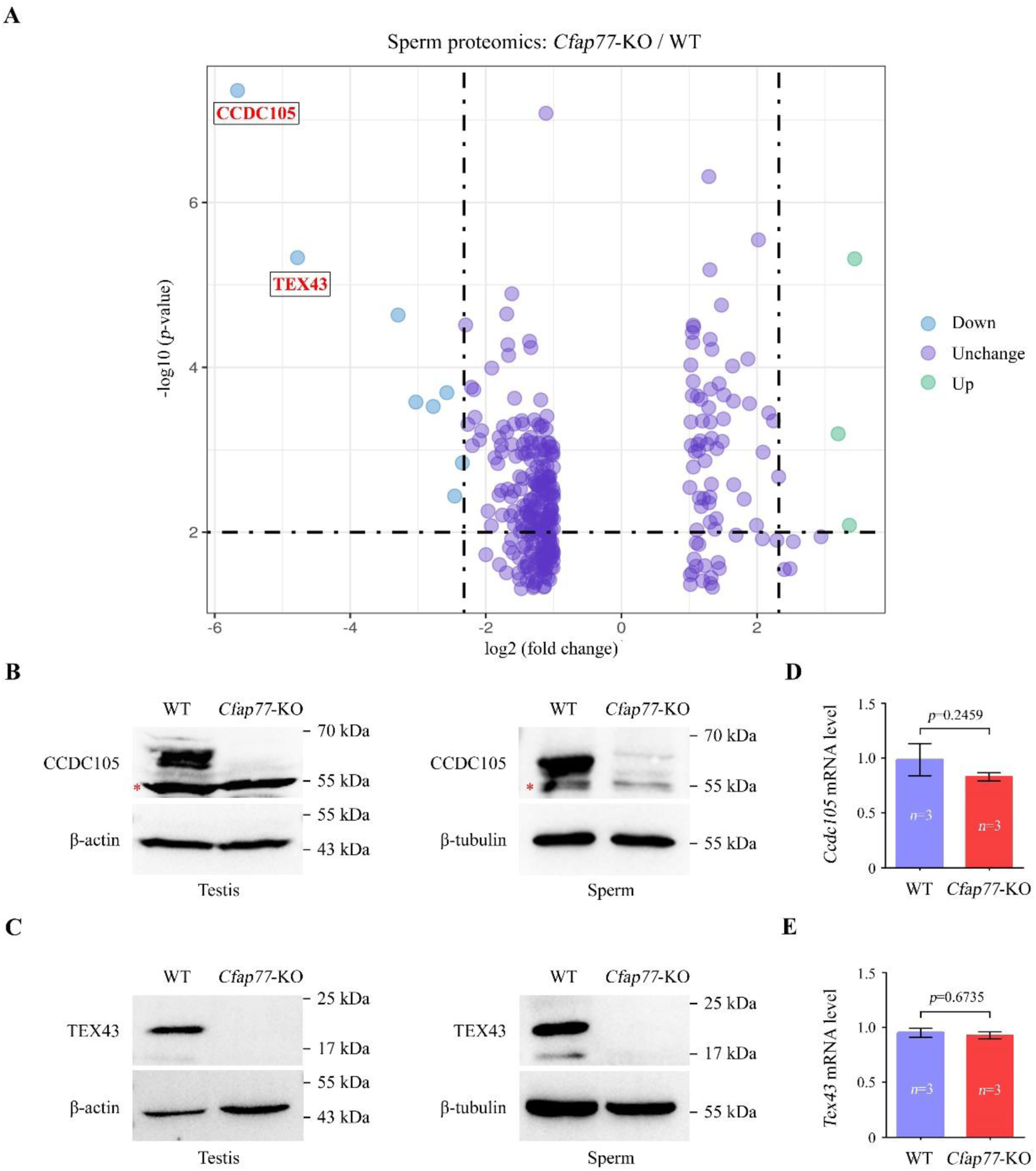
Proteomic analysis of sperm from WT and *Cfap77*-KO mice. **(A)** Quantitative proteomics of sperm protein lysates from WT mice and *Cfap77*-KO mice (*n*=3 each group). Differentially expressed proteins were shown using volcano plot. The X-axis and Y-axis represented fold change (log2) and *p*-value (-log10), respectively. **(B)** Representative immunoblot of CCDC105 in the protein lysates of testes or sperm from WT mice and *Cfap77*-KO mice. β-actin or β-tubulin served as a loading control. Red stars indicated the unspecific band. **(C)** Expression of TEX43 in the protein lysates of testes or sperm from WT mice and *Cfap77*-KO mice. β-actin or β-tubulin served as a loading control. **(D)** Relative mRNA level of *Ccdc105* in testes of WT mice and *Cfap77*-KO mice, as revealed by qRT‒PCR. Student’s *t* test; error bars represent SEM (*n*=3). **(E)** qRT‒PCR result of relative mRNA level of *Tex43* in testes of WT mice and *Cfap77*-KO mice. Student’s *t* test; error bars represent SEM (*n*=3).

### Effect of CFAP77 on other cilia tissues

CFAP77 is a widely conserved OJ protein of axonemal DMTs among eukaryotic ciliated organism^5,7,15,16^, having orthologs showing high similarity in *Chlamydomonas*, mouse, and human (Fig. 6A and Fig. S6). In contrast, CCDC105 and TEX43 are specifically served as CFAP77-interacting proteins in sperm (both sea urchin sperm^16^ and mammalian sperm^7^) (Fig. 6A and Fig. S7). The existing structural models of the CFAP77-CCDC105-TEX43 subcomplex at mouse sperm DMTs, bovine sperm DMTs, and sea urchin sperm DMTs as well as CFAP77 protein at human respiratory DMTs, pig fallopian tube DMTs, pig brain ventricle DMTs, and *Tetrahymena* DMTs were presented in the Fig. S8. Sperm has a single and long flagella and undergoes hyperactivation to fertilize oocyte after a long-distance migration. Thus, CCDC105 and TEX43) are expected to be needed to further strengthen the stability of OJ within axonemal DMTs in sperm.

**Fig 6.**
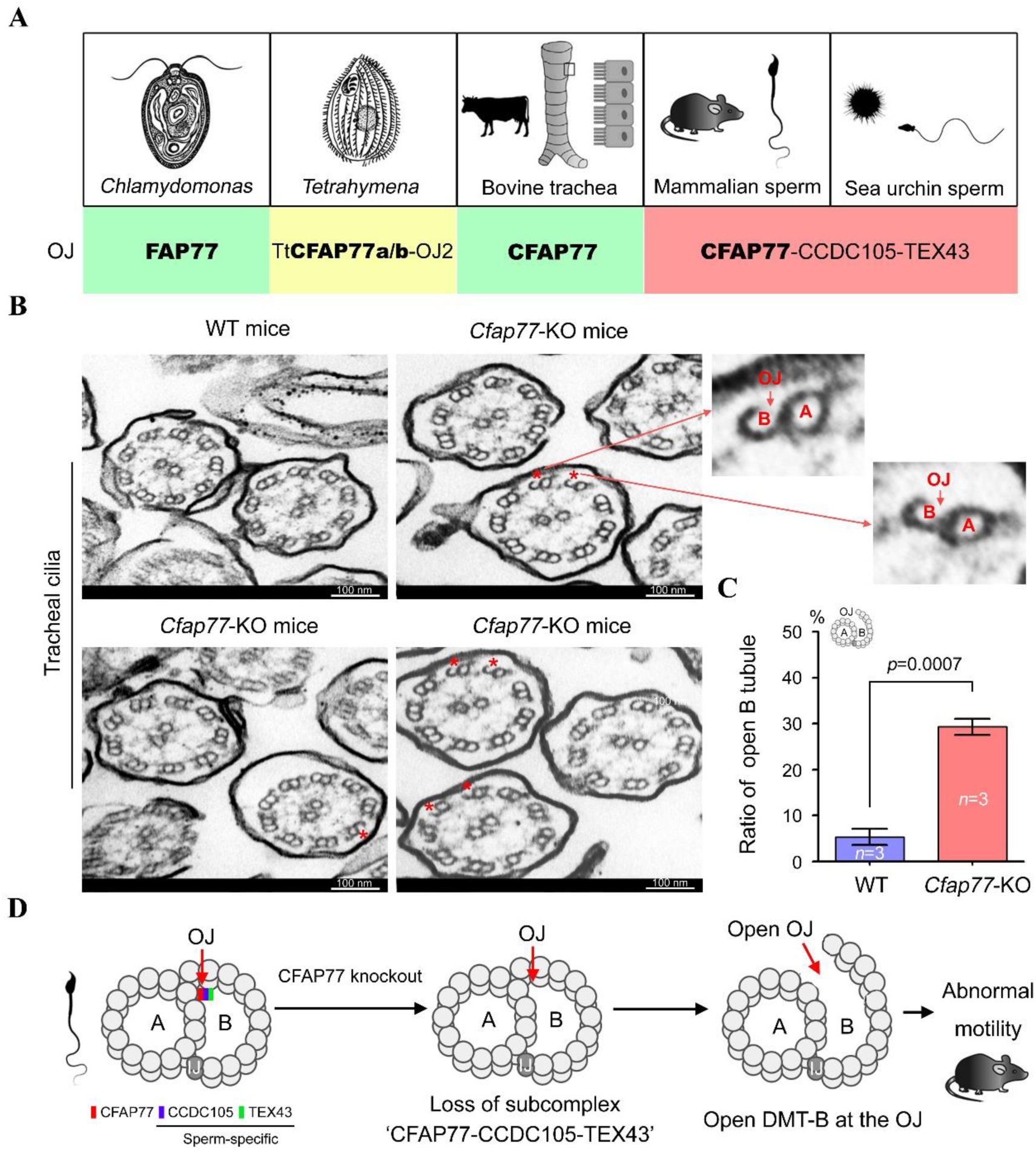
CFAP77, an evolutionary conserved OJ component, is critical for DMT A-B tubules connection of cilia. **(A)** High-resolution cryo-EM/cryo-ET studies indicated that CFAP77 is present among different species, including *Chlamydomonas* (EMD-20631^15^), *Tetrahymena* (EMD-29666^5^), sea urchin (EMD-40619^16^), bovine (EMD-24664^15^), mouse (EMD-35823^7^) and human (EMD-35810^7^). In contrast, CCDC105 and TEX43 are sperm-specific CFAP77-binding proteins. **(B)** TEM analysis of axoneme structures in tracheal cilia from WT mice and *Cfap77*-KO mice. Red stars indicated the OJ region where open DMT-B tubules exists. A single DMT was zoomed in. Scale bars, 100 nm. **(C)** The percentage of axonemes in tracheal cilia with open DMT-B tubules was calculated. Student’s *t* test; error bars represent SEM (*n*=3). **(D)** Schematic diagram to illustrate the connection of DMT-A and B tubules by CFAP77: *Cfap77* knockout leads to the loss of CFAP77-CCDC105-TEX43 subcomplex, and then a disconnection of A-B tubules specifically at the OJ regions, and finally defective motility in mice.

Given that CFAP77 is also expressed in other cilia, we further examined the effect of CFAP77 on ependymal cilia, tracheal cilia and nasal cilia in mice. *Cfap77*-KO mice were healthy with no identifiable abnormalities in cilia function, including laterality abnormalities, hydrocephalus, or respiratory problems. As revealed by scanning electron microscopy (SEM), the morphology of ciliary layer in ependyma, trachea, and nasal cavity was indistinguishable between WT mice and *Cfap77*-KO mice (Fig. S9). Furthermore, the ultrastructures of axonemes in tracheal cilia were examined by the conventional TEM (Fig. 6B). The percentage of axonemes with open DMT-B tubule at the OJ regions in tracheal cilia was significantly increased (5.33%±1.76% in WT mice and 29.33%±1.76% in *Cfap77*-KO mice) after the loss of CFAP77 protein (Fig. 6C). However, this ratio of disconnected A-B tubules of axonemes in tracheal cilia (less than 30%) is lower than that observed in sperm (approximately 45%) (Fig. 3).

Taken all together, our study reveals that knockout of the core OJ protein CFAP77 leads to (i) loss of CFAP77-CCDC105-TEX43 subcomplex in the OJ regions, (ii) and then open DMT-B tubules specifically at the OJ sites, (iii) and finally axoneme instability as well as abnormal sperm motility in mouse sperm (Fig. 6D).

## Discussion

Axoneme, a very large molecular machine, is a cylindrical arrangement of nine DMTs surrounding a CPC. The understanding of axoneme organization is critical for not only cilia biology but also diagnosis of ciliopathies. The DMT-B tubules are incomplete ring of 10 protofilaments and the A and B tubules of each DMT are joined at the OJ and IJ^7^. One of the key questions in axoneme field is how OJ and IJ works to connect A and B tubules within DMTs. High-resolution structure analysis of axonemal DMTs and knockout models provide useful information. CFAP77 is identified to be as the core component of OJ by cryo-EM or *in situ* cryo-ET^5-7,16^ and CFAP77 function is perfectly consistent with its expression. Loss of CFAP77 leads to open DMT-B tubules in *Tetrahymena*^5^ and mouse cilia/flagella (this study), supporting that CFAP77 is an evolutionarily conserved OJ component specifically requiring for DMTs A-B tubule connections in axonemes.

High-resolution information of axonemal ultrastructures is critical to understand their components and assembly^7-9,15,16^. A combination of knockout animal studies with structural biology technologies is one of cutting-edge trends in cilia biology: functional studies are essential to confirm the physiological roles of axonemal proteins and structural biology can further provide in-cell structural insight of how knockout a protein disrupts the axoneme ultrastructure at near-atomic level. A recent study analyzes the sperm DMTs of *Tekt5*-KO mice and compares it with the WT counterpart, providing a first example to apply *in situ* structural technologies to a knockout mouse strain^8^. However, *Tekt5*-KO mice are fertile and sperm motility is normal, prohibiting the biological significance of tektin 5 in axoneme function. Our current study represents a good example to combine structural biology and knockout animals. A loss of CFAP77-CCDC105-TEX43 subcomplex at the OJ region was *in situ* observed in axonemal DMTs of sperm from *Cfap77*-KO mice, representing as the initial even underlying open DMT-B tubules. Moreover, we also found that SPACA9 and ENKUR were missing in the B-tubule and tilted Tektin5 (Tektin5-5) protein was absent in A-tubule lumen of sperm DMTs from *Cfap77*-KO mice. We suggest that the absence of these MIPs in the mutant DMT structure may be the secondary effect of impaired A-B tubule connection.

Previous research on mouse sperm DMTs^7-9,16^ revealed that CCDC105 forms a filament through head-to-tail assembly. TEX43 and CFAP77 further enhance this assembly by interacting with the CCDC105 filaments, reinforcing its linkage with the tubulin wall. Considering a complete loss of CFAP77-CCDC105-TEX43 subcomplex in sperm of *Cfap77*-KO mice, we suggest that the attachment of CCDC105 filaments to the OJ regions may highly dependents on CFAP77. Without CFAP77, CCDC105-TEX43 subcomplex are unable to locate to DMTs and will be degraded in the cytoplasm of spermatids. The protein stability of CCDC105 and TEX43 may be also dependent on CFAP77, but further experiments are needed.

From the perspective of current study, we consider that CFAP77 is the evolutionarily conserved core OJ protein residing between A11-A12 and B01-B02, while CCDC105 and TEX43, two sperm-specific MIPs, may function to joint CFAP77 molecules into a more secure footing for DMTs A-B tubule connection in sperm flagella. It has been recently reported that *Ccdc105*-KO mice exhibit no obvious effect on male reproduction^17^ and TEX43 is involved in the regulation of sperm motility but is also dispensable for male fertility^18^. Given that ultrastructure analyses (e.g., traditional TEM and cryo-EM/cryo-ET) are not performed in sperm axonemes of these knockout mice, whether loss of CCDC105 or TEX43 will partially disturb the DMTs A-B tubule connection is unknown. It is also curious to study the effect of CCDC105 and TEX43 deficiencies on the expression and function of CFAP77. From the viewpoints of the phenotypes of these three knockout mice, the physiological role of CFAP77 on axoneme and sperm motility seems to be much more important than its interacting proteins CCDC105 and TEX43. It is also interesting to generate a *Cfap77* transgenic mouse strain and mate with *Cfap77*-KO mice to study whether CFAP77-CCDC105-TEX43 subcomplex will be rebuilt at the OJ regions and the phenotype of open DMT-B tubules, abnormal sperm motility as well as male infertility will be restored.

CFAP77 is predominantly expressed in the testes, but lower levels of CFAP77 could also be detected in other cilia tissues, including brain, lung, fallopian tube, and retina. Except for male sterility, *Cfap77*-KO mice do not display obvious symptoms of ciliopathy, including laterality abnormalities, hydrocephalus, or defects of respiratory cilia. However, disconnection of A and B tubules within DMTs also happens in cilia cells of *Cfap77*-KO mice. Ultrastructural differences in DMT structures between sperm flagella and motile multiciliated epithelial cells^5,7,15,16^ may account for differential sensitivity to CFAP77 depletion. Furthermore, epithelial cells in ependyma, trachea, and nasal cavity are multiciliated cells and a partial of cilia with open DMT-B tubules may not generate obvious phenotypes. Sperm undergo a long-distance migration with higher oscillation frequency and thus the effect of axoneme instability may be more easily manifested. Indeed, a higher percentage of axoneme exhibits open DMT-B tubules in sperm flagella (approximately 45%) than in other types of cilia (less that 30% in trachea). After sperm hyperactivation, the phenotype of disconnected A-B tubules in sperm axoneme is much more obvious (increasing to ∼70%) in *Cfap77*-KO mice. We also could not exclude the possibility that the effects of CFAP77 on other cilia may exist but more precise inspections are needed.

MIP-variant-associated asthenozoospermia (MIVA) is described as a subtype of asthenozoospermia and is characterized by impaired sperm motility without evident morphological abnormalities^7^. Given that the DMT A-B tubule connections are critical for the stability of sperm axonemes, deficiency in CFAP77-CCDC105-TEX43 subcomplex may be a potential genetic cause of human male infertility with MIVA. Although dynein arms, radial spokes and N-DRC can still attach to the DMTs to form a complete axoneme structure with CPC, the unstable connection of A and B tubules in mutants deficient of this ternary complex will lead to axoneme instability and a cumulating disruption of sperm progressive motility. Indeed, a homozygous mutation in *CCDC105* (c.G1042A/p.E348K) has been identified in male infertile patients with MIVA^7^. However, clinical evidence linking *CFAP77* and *TEX43* mutations to male infertility is still lacking. Whole-exome sequencing of a large cohort of infertile men primarily with MIVA will be useful to screen any pathogenic mutations in *CFAP77* and *TEX43*.

In summary, our study combines *Cfap77*-KO mice and high-resolution *in situ* cryo-ET and reveals that knockout of the core OJ protein CFAP77 leads to loss of CFAP77-CCDC105-TEX43 subcomplex and open DMT-B tubules at the OJ sites, defective sperm motility, as well as male infertility in mammals. Our study not only provides insight into the molecular mechanism of A-B tubule connection within axonemal DMTs but also establishes a novel paradigm to study axoneme by a combination of *in situ* structural biology and gene editing animal models (and possible human patient samples).

## Methods

### Generation of *Cfap77*-KO mice

Animal experiments were approved by the Animal Care and Use Committee of the College of Life Sciences, Beijing Normal University (CLS-AWEC-B-2023-001). The mouse *Cfap77* gene has 5 transcripts and is located on the chromosome 2. Exons 2 and 3 of the *Cfap77-202* (ENSMUST00000157048) transcript were selected as the knockout region. The generation of knockout mice was described in details in our previous studies^19,20^. Briefly, mouse zygotes were coinjected with an RNA mixture of Cas9 mRNA (TriLink BioTechnologies, CA, USA) and sgRNAs. The injected zygotes were transferred into pseudopregnant recipients to obtain the F0 generation. DNA was extracted from tail tissues from 7-day-old offspring and PCR amplification was carried out with genotyping primers (Table S1). A stable F1 generation was obtained by mating positive F0 generation mice with WT C57BL/6JG-pt mice. The gRNA sequence and Sanger sequencing were illustrated in Fig. 2A.

### Generation of polyclonal antibody

The protocol of polyclonal antibody generation was described in details in our previous study^20^. Briefly, recombinant full-length mouse CFAP77 (271 aa) and TEX43 (141 aa) were cloned into the pET-N-His-C-His vector (Beyotime, Shanghai, China) and then transfected into the ER2566 E. coli strain (Weidi Biotechnology, Shanghai, China). Protein expression was induced by 1 mM IPTG at 30°C overnight. After centrifugation, the bacterial pellet was resuspended in buffer (50 mM Tris-HCl pH 8.0, 200 mM NaCl), and the proteins were released by sonication. After centrifugation, anti-His beads (Beyotime) were added to the supernatant and incubated overnight at 4°C. After washing, recombinant protein was eluted with 250 mM imidazole (Beyotime). Recombinant protein was emulsified at a 1:1 ratio (v/v) with Freund’s complete adjuvant (Beyotime) and administered subcutaneously into ICR female mice at multiple points. For the subsequent three immunizations, recombinant protein was emulsified with incomplete Freund’s adjuvant (Beyotime) at an interval of 2 weeks. One week after the last immunization, blood was collected, and the serum was separated.

### Fertility testing

Adult *Cfap77*-KO male mice and their littermate WT mice (*n*=3 each) were mated with WT C57BL/6J females (male: female=1:2) for two months. The vaginal plugs of the mice were examined every morning. Female mice with vaginal plugs were separately fed, and female mice were replenished. The number of pups per litter was recorded.

### In vitro fertilization (IVF)

6-week-old ICR female mice were superovulated by injecting 5 IU of pregnant mare serum gonadotropin (PMSG), followed by 5 IU of human chorionic gonadotropin (hCG) 48 h later. Sperm capacitation was performed for 50 min using TYH solution. Cumulus-oocyte complexes (COCs) were obtained from the ampulla of the uterine tube at 14 h after hCG injection. COCs were then incubated with ∼5 μL sperm suspension in HTF liquid drops at 37°C under 5% CO_2_. After 6 h, eggs were transferred to liquid drops of KSOM medium. Two-cell embryos was counted at 1 day postfertilization. All reagents were purchased from Aibei Biotechnology (Nanjing, China).

### Semen analysis

Sperm counts were determined using a fertility counting chamber (Makler, Israel) under a light microscope. Sperm motility was assessed via the application of a computer-assisted sperm analysis (CASA) system (SAS Medical, China). The sperm suspension was mounted on a glass slide, air-dried, and fixed with 4% PFA for 20 min at room temperature. The slides were stained with Papanicolaou solution (Solarbio, Beijing, China) and observed using a DM500 optical microscope (Leica, Germany).

### Transmission electron microscopy (TEM)

Samples were fixed with 2.5% (vol/vol) glutaraldehyde (GA) in 0.1 M phosphate buffer (PB) at 4°C. Samples were then washed four times in PB and first immersed in 1% (wt/vol) OsO4 and 1.5% (wt/vol) potassium ferricyanide aqueous solution at 4°C for 2 h. After washing, the samples were dehydrated through graded alcohol into pure acetone. Samples were infiltrated in a graded mixture of acetone and SPI-PON812 resin, and then the pure resin was changed. The specimens were embedded in pure resin with 1.5% BDMA, polymerized for 12 h at 45°C and 48 h at 60°C, cut into ultrathin sections (70 nm thick), and then stained with uranyl acetate and lead citrate for subsequent observation and photography with a Tecnai G2 Spirit 120 kV (FEI) electron microscope. All reagents were purchased from Zhongjingkeyi Technology (Beijing, China).

### Western blot

Proteins were extracted using RIPA lysis buffer containing 1 mM PMSF and 2% (v/w) protease inhibitor cocktail (Roche, Basel, Switzerland) on ice. Supernatants were collected following centrifugation at 12,000 × g for 10 min. Proteins were electrophoresed in 10% SDS‒PAGE gels and transferred to PVDF membranes (GE Healthcare, USA). The blots were blocked in 5% milk and incubated with primary antibodies overnight at 4°C, followed by incubation with secondary antibody for 1 h. For primary antibodies, mouse anti-CFAP77 (our homemade, 1:1000), mouse anti-TEX43 (our homemade, 1:1000), and rabbit anti-CCDC105 (Proteintech, 24026-1-AP, 1:1000) were used. Mouse anti-β-Actin (Abcam, ab8226, 1:1000), mouse anti-β-tubulin (Proteintech, a66240-1-Ig, 1:1000), or rabbit anti-Vinculin (Proteintech, 26520-1-AP, 1:1000) served as an internal control. For secondary antibodies, goat anti-rabbit IgG H&L (HRP) (Abmart, M212115, 1:5000) and rabbit anti-mouse IgG H&L (HRP) (Abmart, M212131, 1:5000) was utilized. The signals were evaluated using Super ECL Plus Western Blotting Substrate and a Tanon-5200 Multi chemiluminescence imaging system (China).

### Quantification proteomics

Proteins were extracted from the sperm samples of *Cfap77*-KO mice and WT mice (*n*=3 each group) using 0.1 M Tris-HCL (pH 8.0), 0.1 M dithiothreitol (DTT), 4% SDS, 1 mM PMSF, and 2% (v/w) protease inhibitor cocktail (Roche, Basel, Switzerland), followed by sonication (20% amplitude, 10 pulses, three times) on ice. The supernatants were collected following centrifugation at 12,000 *g* for 20 min. The protocol of mass spectrometry was described in details in our previous studies^20,21^. The mass spectrometry data have been deposited to the ProteomeXchange Consortium via the iProX partner repository with the dataset identifier PXD056128.

### Quantitative RT‒PCR

Total RNA was extracted from the testes of WT mice and *Cfap77*-KO mice using an RNA Easy Fast Tissue/Cell Kit. RNA was converted into cDNA with a FastKing One-Step RT‒PCR Kit according to the manufacturer’s instructions. The cDNAs were used as templates for the subsequent real-time fluorescence quantitative PCR with RealUniversal Colour PreMix (SYBR Green). *Ccdc105* and *Tex43* mRNA expression was quantified according to the 2^−ΔΔCt^. Mouse *Actb* was used as an internal control. Primers were listed in Table S2. All kits were purchased from Tiangen Biotech (Beijing, China).

### Plasmid construction and cell culture

Full-length cDNA encoding CFAP77, CCDC105, and TEX43 was amplified by PCR, cloned into FLAG-, MYC-, or HA-tagged pCMV vectors, and confirmed by Sanger sequencing (Sangon Biotech, Shanghai, China). Mutant plasmids were constructed by using QuickMutation^TM^ Site-Directed Mutagenesis Kit (Beyotime, Shanghai, China). HEK293T cells (ATCC, NY, USA) were cultured at 37°C in a 5% CO_2_ incubator with Dulbecco’s modified Eagle’s medium with 10% fetal bovine serum and 1% penicillin‒ streptomycin (Gibco, NY, USA).

### Coimmunoprecipitation (co-IP)

HEK293T cells were transfected with FLAG-, MYC-, and/or HA-tagged plasmids by Lipofectamine 3000 (Thermo Fisher, CA, USA). 48 hours after transfection, cells were lysed with Pierce™ IP Lysis Buffer (Thermo Fisher) containing a 2% (v/w) protease inhibitor cocktail (Roche, Basel, Switzerland) for 30 min at 4°C and then centrifuged at 12,000 ×g for 10 min. Protein lysates were incubated overnight with FLAG antibody (Abmart, PA9020, 2 μg) at 4°C. The lysates were then incubated with 20 μL Pierce™ Protein A/G-conjugated Agarose for 4 h at 4°C. The agarose beads were washed five times with Pierce™ IP Lysis Buffer and boiled for 5 min in 1×SDS loading buffer. Input and IP samples were analysed by Western blotting using HRP conjugated anti-FLAG antibody (Abmart, PA9020, 1:1000), anti-MYC antibody (Abmart, M20019, 1:1000), or anti-HA antibody (Abmart, M20003, 1:1000).

### Sample preparation of mouse sperm axoneme

Freshly extracted sperm were centrifuged at 400 G (Thermo Scientific Legend Micro 17 R) for 5 min at 4°C. The precipitate per 100 μL semen was carefully re-suspended in 100 μL of precooled PBS and diluted 5.5-fold with PBS before use. The cryo-EM grid (Quantifoil R2/1, Au 200 mesh) was discharged for 60 s using Gatan Solarus. Sperm samples were quickly frozen by vitrification using Leica EMGP. Samples diluted in 3 μL PBS were immediately water absorbed at 100% relative humidity and 4°C for 4s, and then the frozen samples were placed in a mixture of ethane and methane cooled to -195°C and stored in liquid nitrogen for cryo-FIB thinning. The cryo-FIB thinning strategy was referred to in our previous work^9^.

### Cryo-ET tilt series collection

The grid after the reduction of cryo-FIB was mounted onto Autoloader in Titan Krios G3 (Thermofisher Scientific) 300 KV TEM, equipped with a Gatan K2 direct electron detector (DED) and a BioQuantum energy filter. Tilt series were collected at a magnification of ×42,000, resulting in a physical pixel size of 3.4 Å in K2 DED and 3.4 Å in counting mode. Before data collection, the pre-tilt of the sample was determined visually, and the pre-tilt was set to 10°or -9°to match the pre-determined geometry induced by loading grids. The total dose for each tilt was set to 3.5 electrons per angstrom squared, divided into 10 frames over a 1.2-s exposure, and the tilt angle was set to -9°pre-tilt of -66°to +51°, or +10°pre-tilt of -50°to +67°, in steps of 3°, resulting in 40 tilts and 140 electrons per tilt series. The slit width was set to 20 eV, zero loss peaks were refined after the collection of each tilt series, and nominal defocus was set from -1.8 to -2.5 μm. All tilt series used in this study were collected using a beam-image-shift facilitated acquisition scheme based on a dose symmetry strategy using a script developed in-house in SerialEM software^22-24^.

### Data processing

After collecting data, all fractioned movies were imported into Warp for essential processing, including motion correction, Fourier binning by a factor of 1 of the counting mode frames, CTF estimation, masking platinum islands or other high-contrast features, and tilt series generation^25^. Subsequently, AreTomo is used to automatically align the tilt series^25,26^. Conduct a visual inspection of the aligned tilt series in IMOD and delete any low-quality frames (such as those blocked by sample tables or grid bars, containing visible crystalline ice or showing significant jumps) to create the new tilt series in Warp. The new tilt series has undergone a second round of AreTomo alignment. Then, using the same criteria as in the first round, low-quality frames are again deleted. The new tilt series then undergoes a third round of automatic alignment, continuing the process until no frames need to be removed. After the alignment of the tilt series, those tilt series that are less than 30 frames or fail are not further processed^26,27^. All remaining alignment parameters for the tilt series were passed back to Warp, and an initial layer map reconstruction was performed in Warp with a pixel size of 27.2 Å. Among the total 174 tomograms, we used the filament picking tool in Dynamo to manually pick DMT particles from 100 of all tomograms. By selecting the starting and ending points of each DMT fiber, and separating each cutting point by 8 nm along the fiber axis, we were able to obtain 100 sets of DMT particles^28^. The 3D coordinates and two of the three Euler angles (except for in-plane rotation) are automatically generated by Dynamo and then transferred back to Warp for exporting sub-tomograms^25,28,29^. In RELION 3.1, sub-tomograms were refined, using the ABTT package to transform the RELION star file and Dynamo table file and use Dynamo and/or RELION to jointly generate a mask^30,31^. First, all particles were reconstructed into the box size of 48^3^ voxels with a pixel size of 27.2 Å, and all extracted particles were directly averaged and low-pass filtered at 80 Å to generate an initial reference. Then 3D classification with K = 1 was performed under the constraints of the first two Euler angles (—sigma_tilt 3 and —sigma_psi 3 in RELION), and 3D automatic refinement was performed after 25 iterations. After alignment, manually cleaned the particles in ChimeraX-1.6^32^, and then transferred these aligned parameters back to Warp to export the sub-tomograms with a box size of 843 voxels and a pixel size of 13.6 Å. The particles were automatically refined in RELION. Then transferred these aligned parameters back to Warp to export the sub-tomograms with the box size of 128^3^ voxels and the pixel size of 6.8 Å. Then it was automatically refined in RELION. After removing any duplicate particles in Dynamo, it was automatically refined in M with a final resolution of 24 Å^33^. Based on the previous atomic model of DMT in WT mice (PDB: 8I7O), we deleted the models of CFAP77, CCDC105, and TEX43 in ChimeraX-1.6 to obtain the DMT model of *Cfap77*-KO mouse sperm^9^.

### Statistical analyses

Data are presented as the mean ± standard error (SEM) and were analysed using GraphPad Prism version 5.01 (GraphPad Software). Student’s *t* test (unpaired, two-tailed) was used for the statistical analyses.

## Acknowledgments

We would like to thank Center for Biological Imaging (CBI), Institute of Biophysics (IBP), Chinese Academy of Science (CAS) for the support of TEM, specimen vitrification, cryo-FIB milling, and cryo-ET data collection. We also acknowledge Jin Liu and Chao Xi from the Experimental Technology Center for Life Sciences, Beijing Normal University, for providing experimental platform.

## Author contributions

SC and YZ conceived and designed the project, BW designed animal model and animal experiments, LX performed the mouse experiments, GY performed the cryo-ET experiments, YL performed the molecule experiments, SC wrote the manuscript, FS and YZ revised the manuscript.

## Data availability

Proteomics data have been deposited to the ProteomeXchange Consortium via the iProX partner repository with the dataset identifier PXD056128. The map of sperm DMTs from *Cfap77*-KO mice has been deposited in the EMDB under accession codes EMD-63176.

## References

1. Teves, M.E., Nagarkatti-Gude, D.R., Zhang, Z. & Strass, JF.3rd. Mammalian axoneme central pair complex proteins: Broader roles revealed by gene knockout phenotypes. Cytoskeleton.73,3–22 (2016).

2. Klena, N. & Pigino, G. Structural biology of cilia and intraflagellar transport. Annu Rev Cell Dev Biol. 38,103–123 (2022).

3. Jiao, S.Y., Yang, Y.H. & Chen, S.R. Molecular genetics of infertility: loss-of-function mutations in humans and corresponding knockout/mutated mice. Hum Reprod Update. 27,154–189 (2021).

4. Khalifa, A.A.Z. et al. The inner junction complex of the cilia is an interaction hub that involves tubulin post-translational modifications. Elife. 9:e52760 (2020).

5. Kubo, S. et al. Native doublet microtubules from Tetrahymena thermophila reveal the importance of outer junction proteins. Nat Commun. 14,2168 (2023).

6. Gui, M. et al. SPACA9 is a lumenal protein of human ciliary singlet and doublet microtubules. Proc Natl Acad Sci U S A.119,e2207605119 (2022).

7. Zhou, L. et al. Structures of sperm flagellar doublet microtubules expand the genetic spectrum of male infertility. Cell. 186,2897–2910 (2023).

8. Chen, Z. et al. De novo protein identification in mammalian sperm using in situ cryoelectron tomography and AlphaFold2 docking. Cell. 186,5041–5053. (2023).

9. Tai, L., Yin, G., Huang, X., Sun, F. & Zhu, Y. In-cell structural insight into the stability of sperm microtubule doublet. Cell Discov. 9,116 (2023).

10. Dymek, E.E. et al. PACRG and FAP20 form the inner junction of axonemal doublet microtubules and regulate ciliary motility. Mol Biol Cell. 30,1805–1816 (2019).

11. Chen, Z., Li, M., Zhu, H. & Ou, G. Modulation of inner junction proteins contributes to axoneme differentiation. Proc Natl Acad Sci U S A. 120,e2303955120 (2023).

12. Chrystal, P.W. et al. The inner junction protein CFAP20 functions in motile and non-motile cilia and is critical for vision. Nat Commun. 13,6595 (2022).

13. Liu, C. et al. Deleterious variants in X-linked CFAP47 induce asthenoteratozoospermia and primary male infertility. Am J Hum Genet. 108,309–323 (2021).

14. Zhu, Y. et al. In-cell structural insight into the asymmetric assembly of central apparatus in mammalian sperm axoneme. bioRxiv (2024). 10.1101/2024.08.06.606614.

15. Gui, M. et al. De novo identification of mammalian ciliary motility proteins using cryo-EM. Cell. 184,5791–5806 (2021).

16. Leung, M.R. et al., Structural specializations of the sperm tail. Cell. 186,2880–2896 (2023).

17. Nguyen, T.T.T. et al. Gene-deficient mouse model established by CRISPR/Cas9 system reveals 15 reproductive organ-enriched genes dispensable for male fertility. Front Cell Dev Biol. 12,1411162 (2024).

18. Miyata, H. et al. SPATA33 localizes calcineurin to the mitochondria and regulates sperm motility in mice. Proc Natl Acad Sci U S A. 118,e2106673118 (2021).

19. Zhang, X.Z., Wei, L.L., Jin, H.J., Zhang, X.H. & Chen, S.R. The perinuclear theca protein Calicin helps shape the sperm head and maintain the nuclear structure in mice. Cell Rep. 40,111049 (2022).

20. Geng, X.Y., Jin, H.J., Xia, L., Wang, B.B. & Chen, S.R. Tektin bundle interacting protein, TEKTIP1, functions to stabilize the tektin bundle and axoneme in mouse sperm flagella. *Cell Mol Life Sci*. **81**,118 (2024).

21. Jin, H.J. et al. CFAP70 is a solid and valuable target for the genetic diagnosis of oligo-astheno-teratozoospermia in infertile men. EBioMedicine. 93,104675 (2023).

22. Hagen, W.J.H., Wan, W. & Briggs, J.A.G. Implementation of a cryo-electron tomography tilt-scheme optimized for high resolution subtomogram averaging. J Struct Biol. 197,191–198 (2017).

23. Wu, C., Huang, X., Cheng, J., Zhu, D. & Zhang, X. High-quality, high-throughput cryo-electron microscopy data collection via beam tilt and astigmatism-free beam-image shift. J Struct Biol. 208,107396 (2019).

24. Mastronarde, D.N. Automated electron microscope tomography using robust prediction of specimen movements. J Struct Biol. 152,36–51 (2005).

25. Tegunov, D. & Cramer, P. Real-time cryo-electron microscopy data preprocessing with Warp. Nat Methods. 16,1146–1152 (2019).

26. Zheng, S. AreTomo: An integrated software package for automated marker-free, motion-corrected cryo-electron tomographic alignment and reconstruction. J Struct Biol X. 6,100068 (2022).

27. Kremer, J.R., Mastronarde, D.N. & McIntosh, J.R. Computer visualization of three-dimensional image data using IMOD. J Struct Biol.116,71–6 (1996).

28. Castaño-Díez, D., Kudryashev, M., Arheit, M. & Stahlberg, H. Dynamo: a flexible, user-friendly development tool for subtomogram averaging of cryo-EM data in high-performance computing environments. J Struct Biol. 178,139–51 (2012).

29. Burt, A., Gaifas, L., Dendooven, T. & Gutsche, I. A flexible framework for multi-particle refinement in cryo-electron tomography. PLoS Biol. 19,e3001319 (2021).

30. Zivanov, J. et al. New tools for automated high-resolution cryo-EM structure determination in RELION-3. Elife. 7:e42166 (2018).

31. Scheres, S.H.W. RELION: implementation of a Bayesian approach to cryo-EM structure determination. J Struct Biol. 180,519–30 (2012).

32. Pettersen, E.F. et al. UCSF Chimera--a visualization system for exploratory research and analysis. J Comput Chem. 25,1605–12 (2004).

33. Tegunov, D., Xue, L., Dienemann, C., Cramer, P. & Mahamid, J. Multi-particle cryo-EM refinement with M visualizes ribosome-antibiotic complex at 3.5 Å in cells. Nat Methods.18,186–193 (2021).

