## Supplemental Files for "Essential role of CFAP77-CCDC105-TEX43 subcomplex in connecting axonemal A and B tubules for mammalian sperm motility"

**The PDF file includes:**

Methods
Fig. S1 to S9
Tables S1 to S2
References

**Other Supplementary Materials for this manuscript include the following:**

Movies S1 to S2

### Methods

#### Histological analysis

The testes and caudal epididymis were dissected and fixed in 4% PFA overnight at 4 ℃. Fixed tissues were embedded in paraffin, sectioned (5 μm thick), dewaxed, and rehydrated. The sections were stained with periodic acid schiff (PAS) solution or hematoxylin-eosin (H&E) solution (Solarbio, Beijing, China) before imaging using a Leica DM-500 optical microscope (Leica Microsystems, German).

#### Peanut agglutinin (PNA) staining

After permeabilization with 1% Triton X-100 for 1 h, the slides of mouse testis section were stained for PNA-TRITC dye (Sigma-Aldrich, CA, USA) for 1 h. After washing, the slides were counterstained with DAPI dye (Solarbio) and imaged with a fluorescence microscope (Leica).

#### Scanning electron microscope (SEM)

Tissues were fixed in 2.5% phosphate-buffered glutaraldehyde (GA) (Zhongjingkeyi Technology, Beijing, China) at room temperature for 30 min and then deposited on coverslips. The coverslips were dehydrated via an ascending gradient of 50%, 70%, 95%, and 100% ethanol and air-dried. Specimens were then attached to specimen holders and coated with gold particles using an ion sputter coater before being viewed with a JSM-IT300 scanning electron microscope (JEOL, Tokyo, Japan).

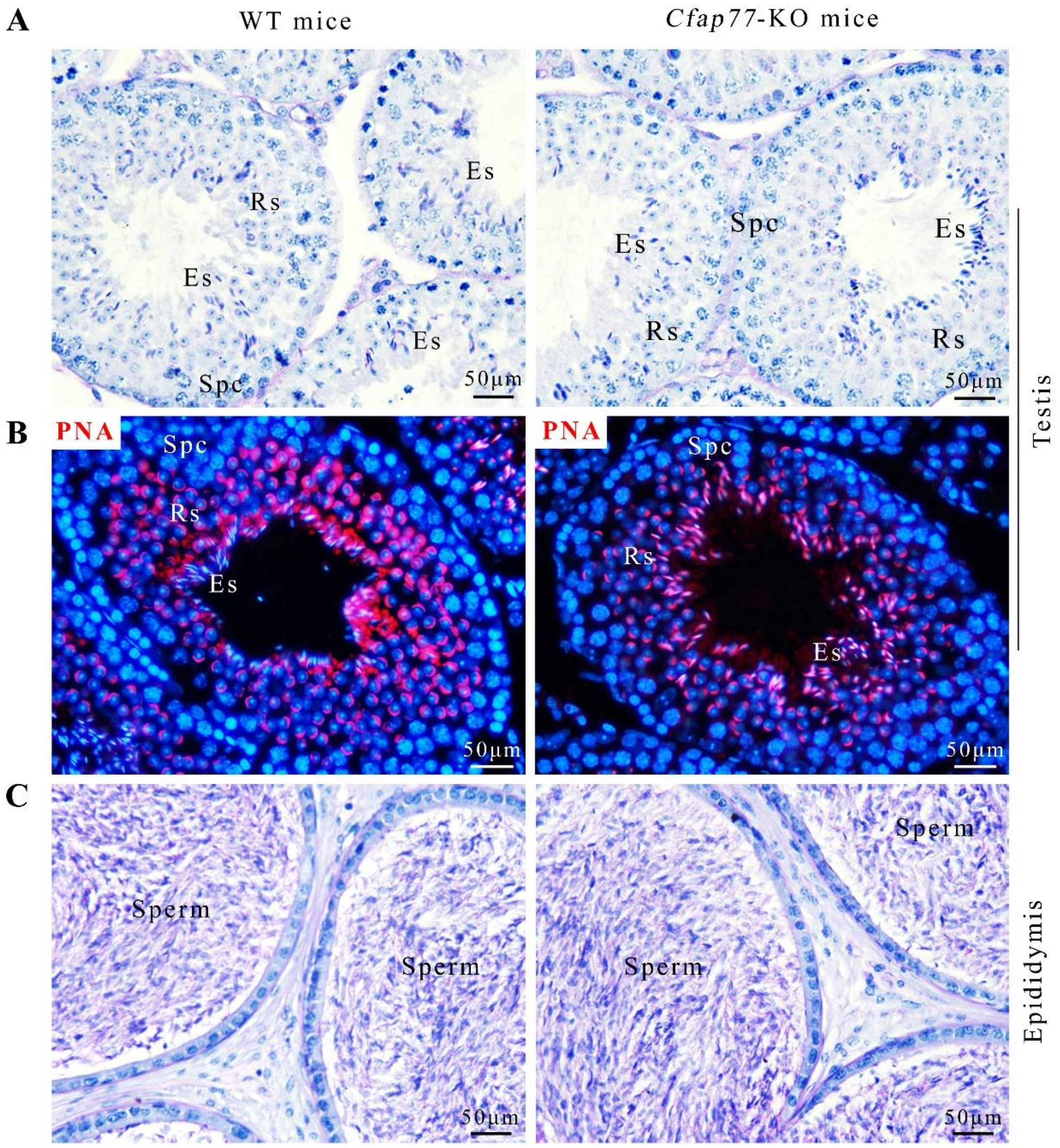


**Fig S1. Analysis of the reproductive phenotype of *Cfap77*-KO mice.**

**(A)** Representative histological morphology of testis sections from *Cfap77*-KO mice and WT mice by Periodic Acid Schiff staining. Spc, spermatocytes, Rs, round spermatids, Es, elongating spermatids. Scale bars, 50 μm.

**(B)** Staining of PNA-TRITC to reveal the acrosomal formation using testis section from *Cfap77*-KO mice and WT mice. Nuclei was stained with DAPI.

**(C)** Representative histological morphology of the cauda epididymis from *Cfap77*-KO mice and WT mice by H&E staining.

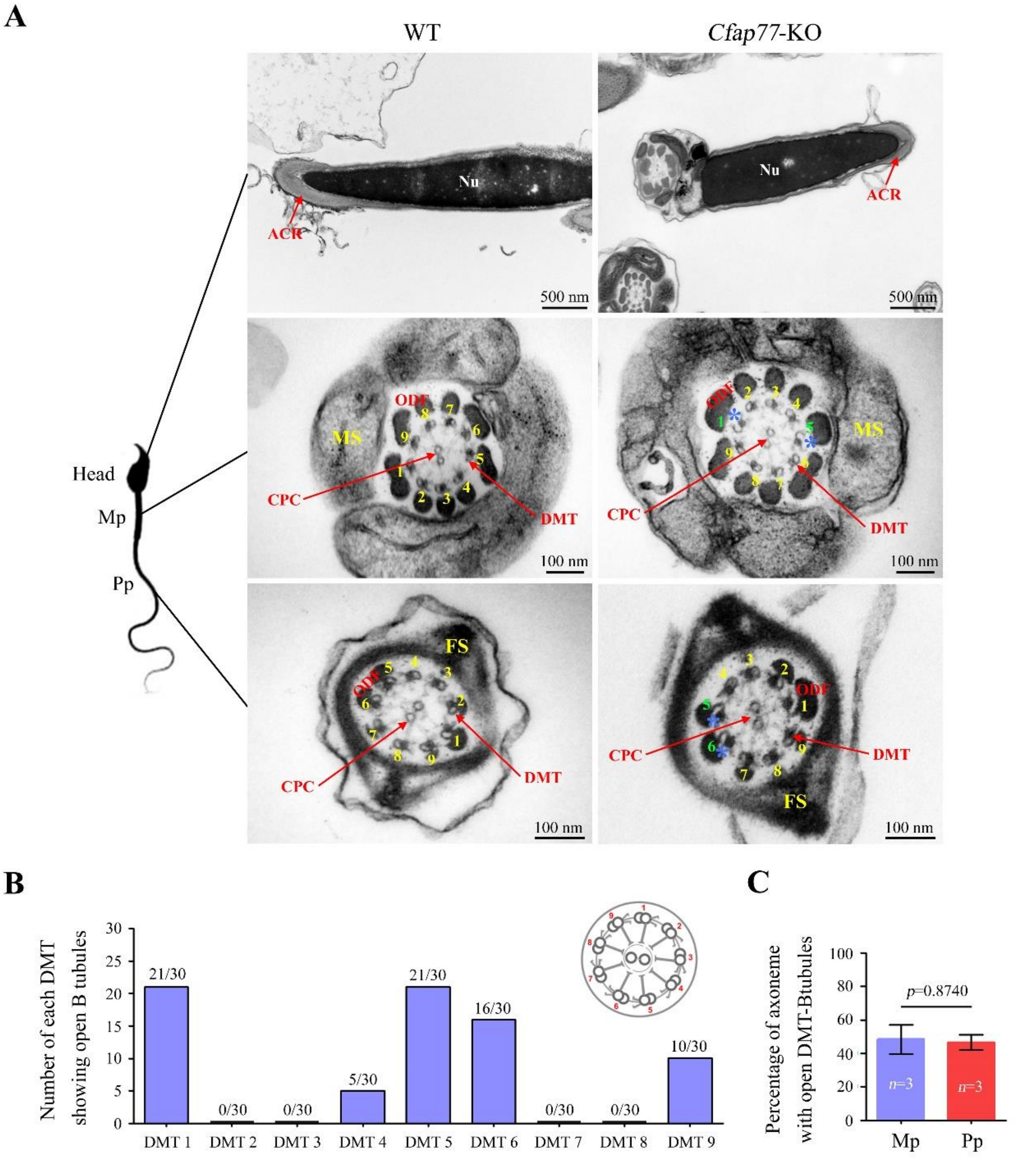


**Fig S2. Detailed analysis of opened DMT-B tubules of sperm flagella in *Cfap77*-KO mice.**

**(A)** TEM analysis of acrosome (ACR), mid-piece (Mp) and principal piece (Pp) of tails in WT mice and *Cfap77*-KO mice. Nu, nucleus; MS, mitochondrial sheath; FS, fibrous sheath; CPC, central pair complex; DMT, doublet microtubule; ODF, outer dense fiber. Scale bars, 100 nm or 500 nm.

**(B)** Number of each DMT (1-9) exhibiting the open B tubules. A total of 30 axonemes with open DMT-B tubules in *Cfap77*-KO sperm were counted.

**(C)** Percentage of axonemes with open DMT-B tubules was calculated between Mp and Pp of sperm flagella in *Cfap77*-KO mice. Student's *t* test; error bars represent SEM ($n$=3).

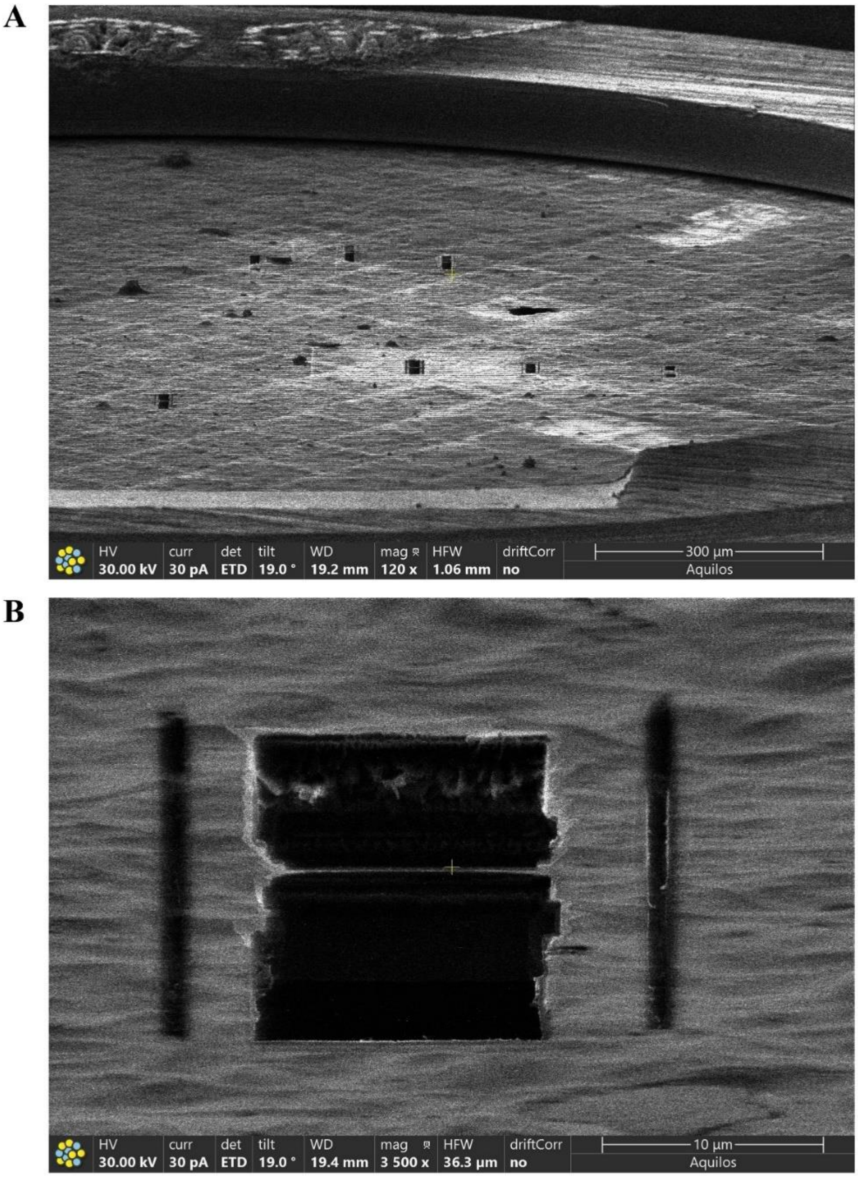


**Fig S3. Cryo-FIB milling of sperm from *Cfap77*-KO mice.**
**(A)** Inspection of frozen sperm on the grid.
**(B)** Inspection of frozen sperm on the grid after FIB milling. The thin lamellae were used for data collection.

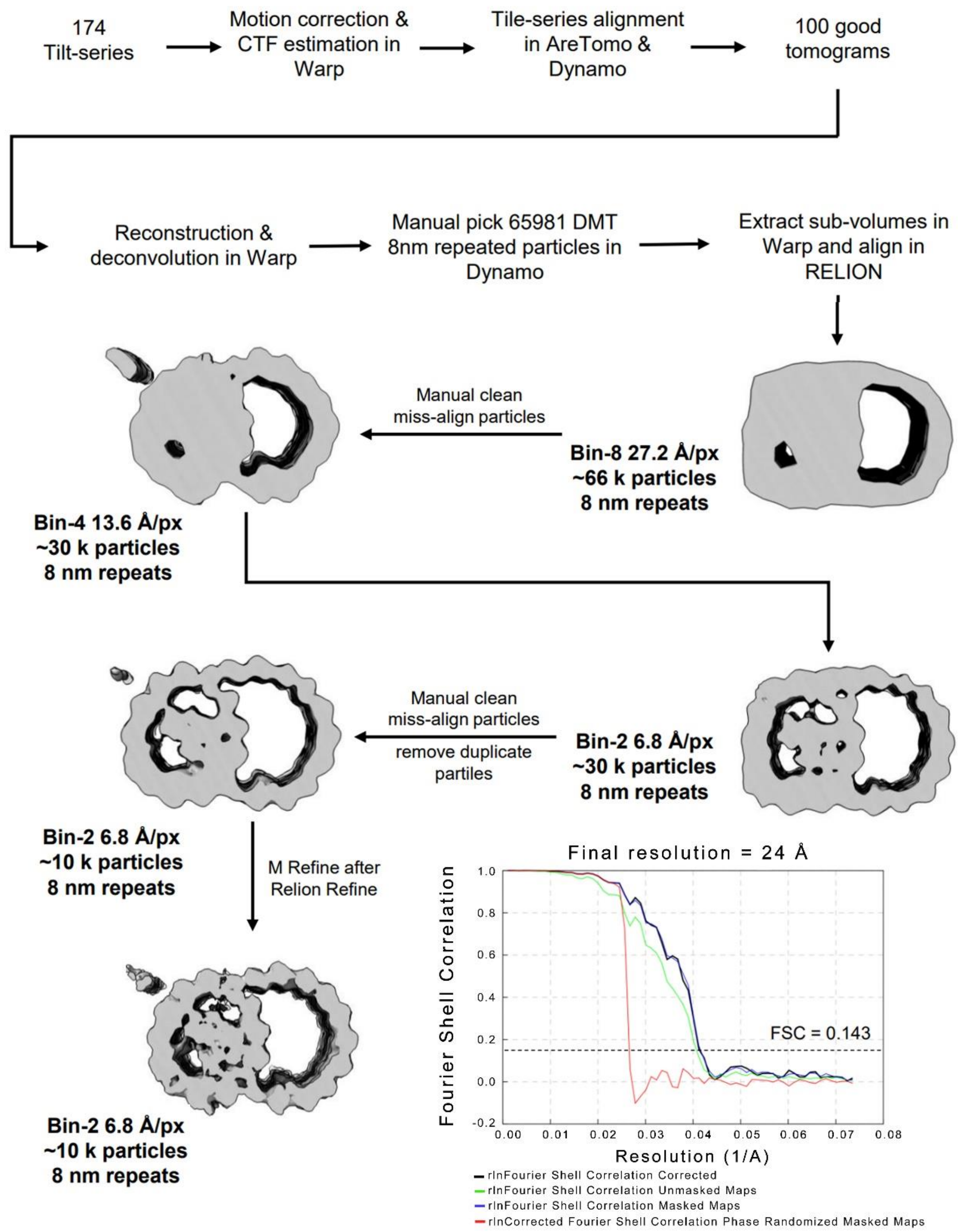


**Fig S4. The data processing of sperm DMTs from *Cfap77*-KO mice**. The pixel sizes at different binning levels are indicated in angstroms per pixel (Å/px for short). The half-map Fourier Shell Correlation (FSC) plot is shown.

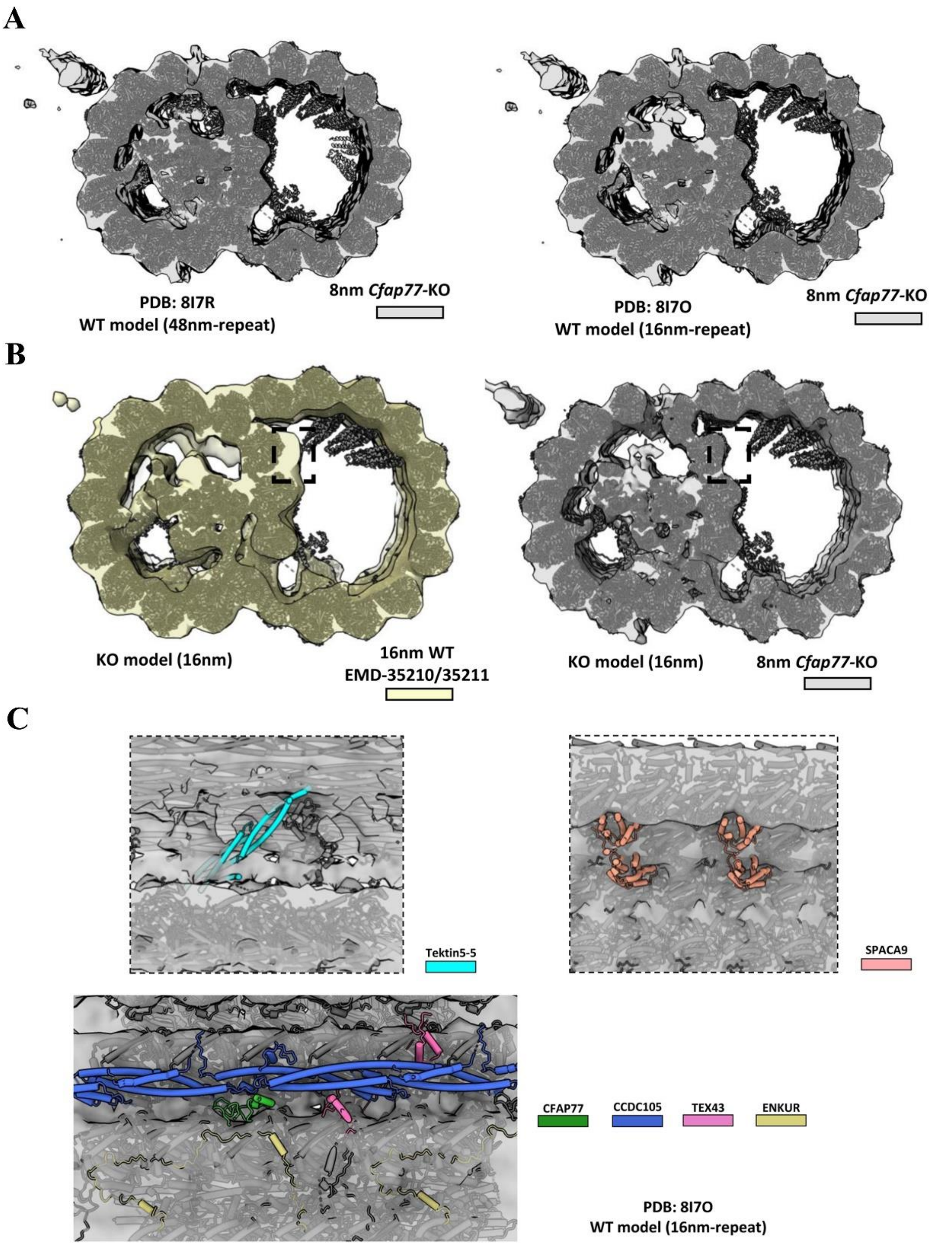


**Fig S5. Structural comparison of sperm DMTs from WT and *Cfap77*-KO mice.**
**(A)** The 48 nm repeat (PDB: 8I7R) and 16 nm repeat (PDB: 8I7O) WT DMT models were fitted in the DMT map of *Cfap77*-KO mice.
**(B)** The model of *Cfap77*-KO DMTs was fitted in the 16 nm repeat DMT maps of WT mice and *Cfap77*-KO mice. Dashed box indicated the difference density near protofilaments A11 and A12.
**(C)** The atomic model of 16 nm-repeat WT DMTs (PDB: 8I7O) fitted into the density map of *Cfap77*-KO DMTs. The lost components of Tektin5-5, SPACA9, CFAP77, CCDC105, TEX43, and ENKUR are indicated.

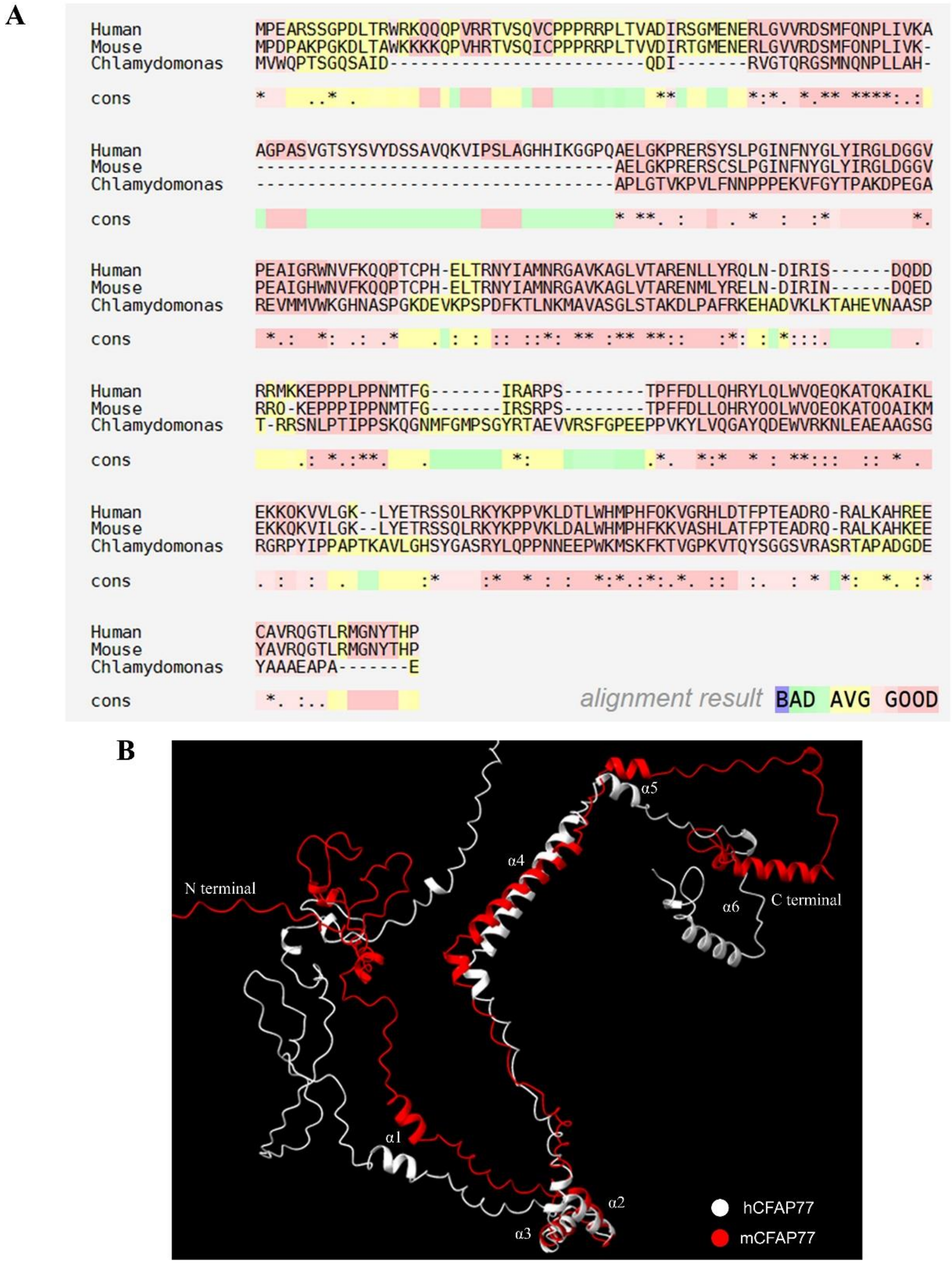


**Fig S6. CFAP77 is conserved among *Chlamydomonas*, mouse, and human.**

**(A)** Using sequence alignment analysis (M-Coffee), we revealed that CFAP77 was remarkably conserved among *Chlamydomonas*, mouse, and human.

**(B)** Alignment analysis of three-dimensional structures of mouse CFAP77 and human CFAP77. Their structures were predicted by AlphaFold 3.

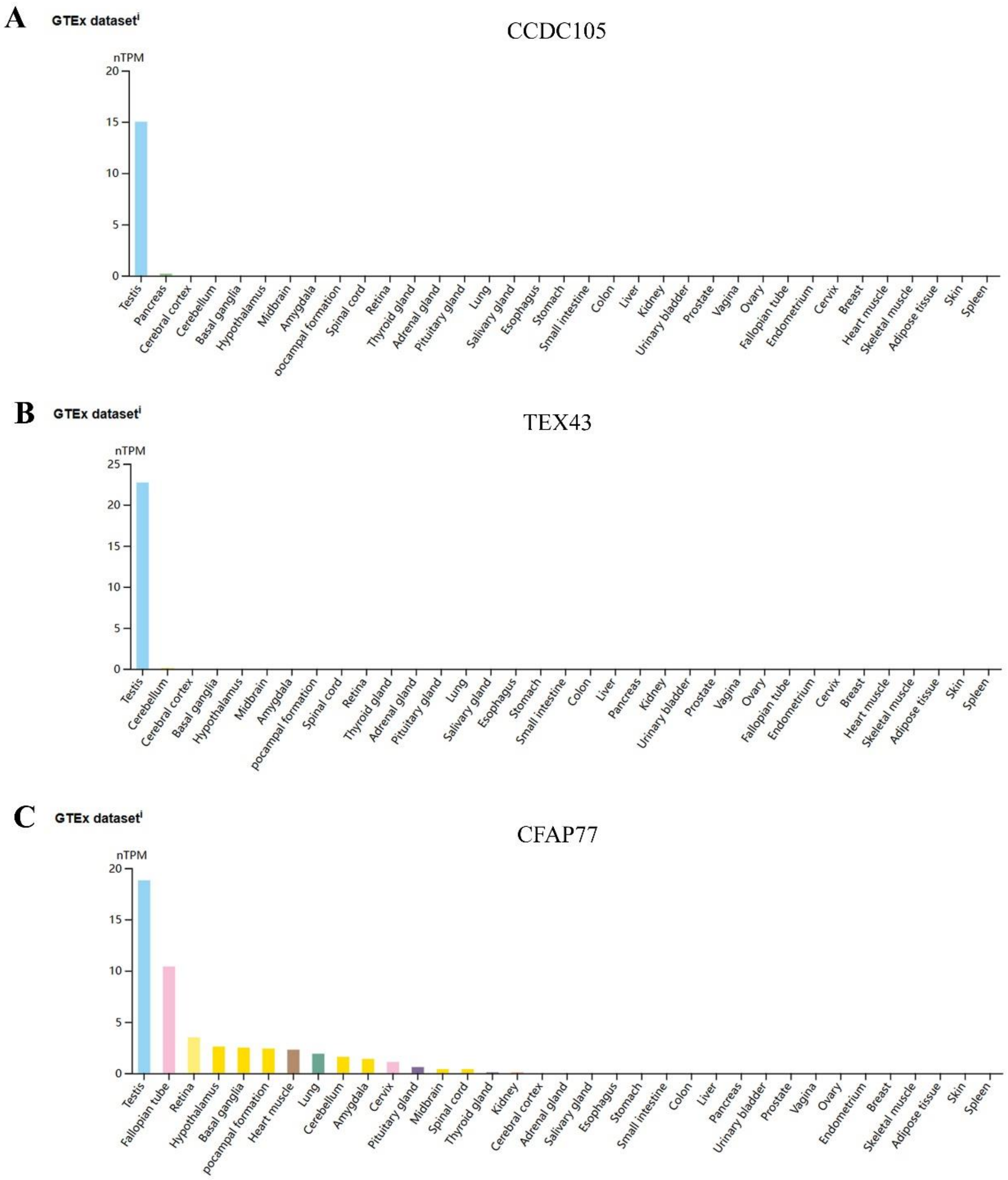


**Fig S7. Expression information of CFAP77, CCDC105, and TEX43 in humans.**

**(A)** According to the Human Protein Atlas (HPA) database, *CCDC105* mRNA was restricted to testes.

**(B)** *TEX43* mRNA was also restricted to testes.

**(C)** Among different tissues, *CFAP77* mRNA was predominantly expressed in testes, but also present in other cilia tissues, including fallopian tube, retina, brain, and lung.

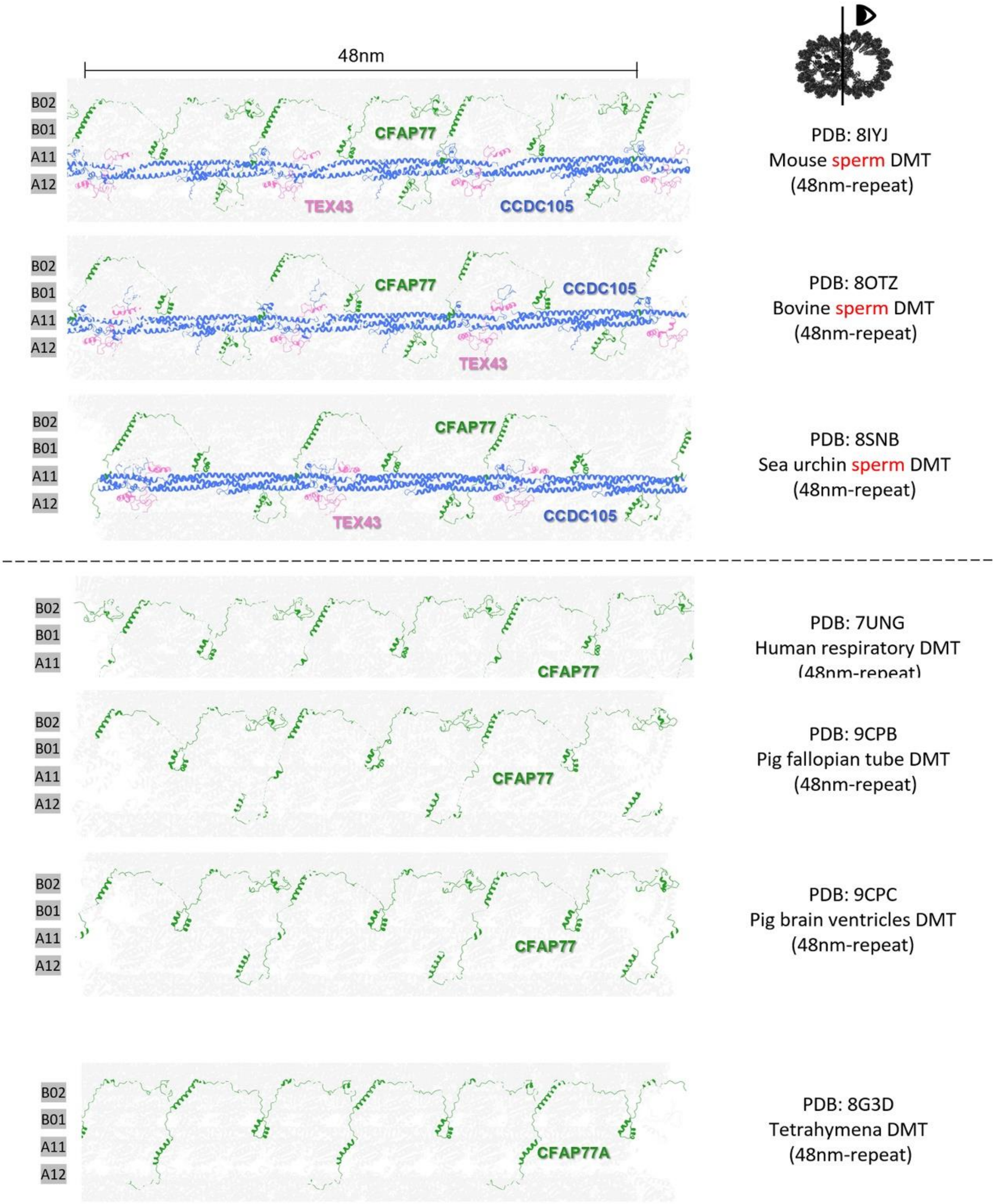


**Fig S8. Cryo-EM/cryo-ET structural models of CFAP77, CCDC105, TEX43.** Structural models of the CFAP77-CCDC105-TEX43 subcomplex or CFAP77 at the OJ regions of axonemes from various species and tissues are presented, with a vertical section showing a 48 nm length of the DMT. From top to bottom, the models include: mouse sperm (PDB: 8IYJ), bovine sperm (PDB: 8OTZ), sea urchin sperm (PDB: 8SNB), human respiratory epithelium (PDB: 7UNG), pig fallopian tube (PDB: 9CPB), pig brain ventricles (PDB: 9CPC), and Tetrahymena (PDB: 8G3D).

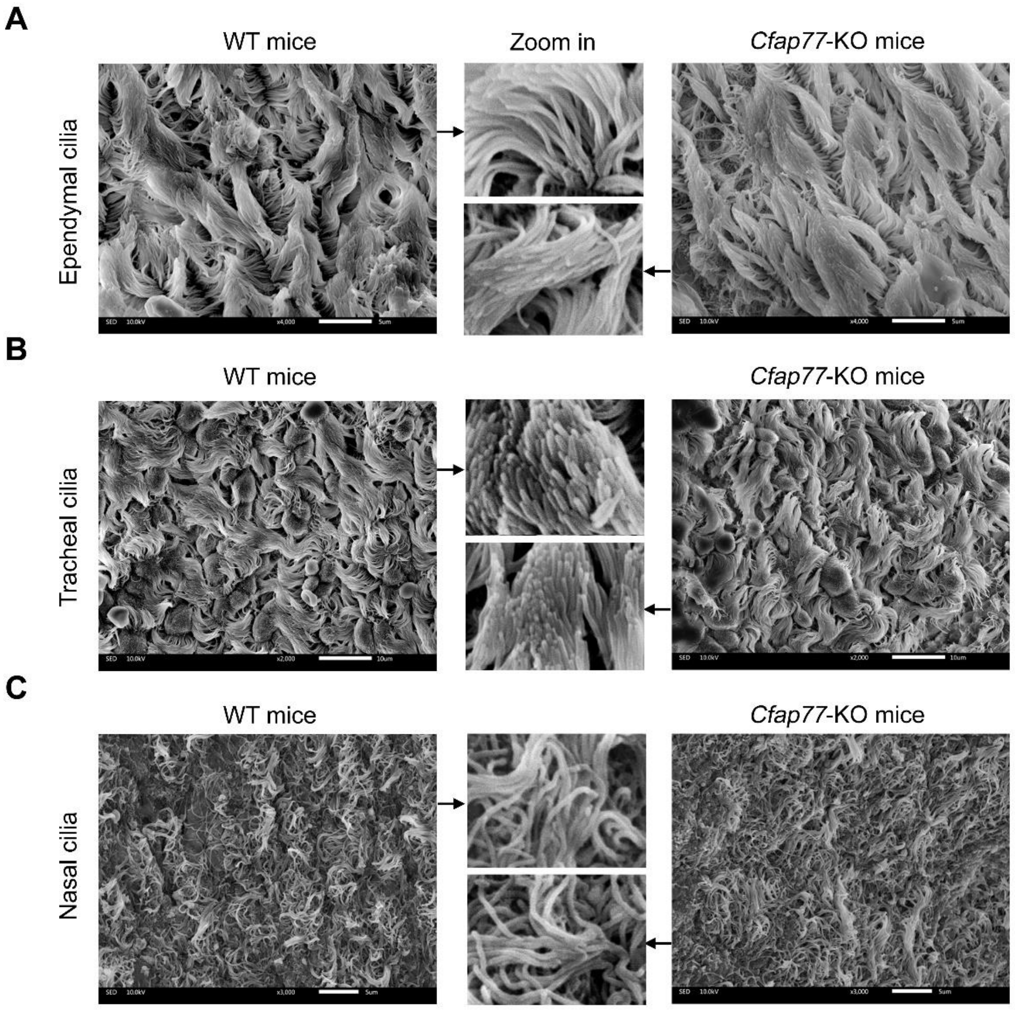


**Fig S9. SEM analysis of cilia in *Cfap77*-KO mice.**

**(A)** Representative SEM images of brain ependymal cilia from *Cfap70*-KO mice and WT mice. Scale bar, 5 μm.

**(B)** Representative SEM images of tracheal cilia from *Cfap70*-KO mice and WT mice. Scale bar, 10 μm.

**(C)** Representative SEM images of nasal cilia from *Cfap70*-KO mice and WT mice. Scale bar, 5 μm.

**Table S1. Primers for mouse genotyping.**

| Primers | Sequence | Size |
|---|---|---|
| F1 | 5’-GTCTCGGTTGCTCATTCCTATGT-3’ | Targeted: 484 bp |
| R1 | 5’-GTGTCCCTCATGACCTTCAAGAAA-3’ | |
| F1 | 5’-GTCTCGGTTGCTCATTCCTATGT-3’ | WT: 827 bp |
| R2 | 5’-CTGTGCCATTTGACTATGGCCTAC-3’ | |

**Table S2. Primers for qRT-PCR.**

| Target | Sequence | Size |
|---|---|---|
| *Ccdc105* | F: TACCGACCTAAGTGTGAGAAGATCC | 195 bp |
| | R: CATGAGTTCCACAGCTTGCAG | |
| *Tex43* | F: GAACGTGAATCTACAGCAAGCA | 115 bp |
| | R: GACGGCTTTCCGATCTTCCC | |
| *Actb* | F: AACAGTCCGCCTAGAAGCAC | 281 bp |
| | R: CGTTGACATCCGTAAAGACC | |

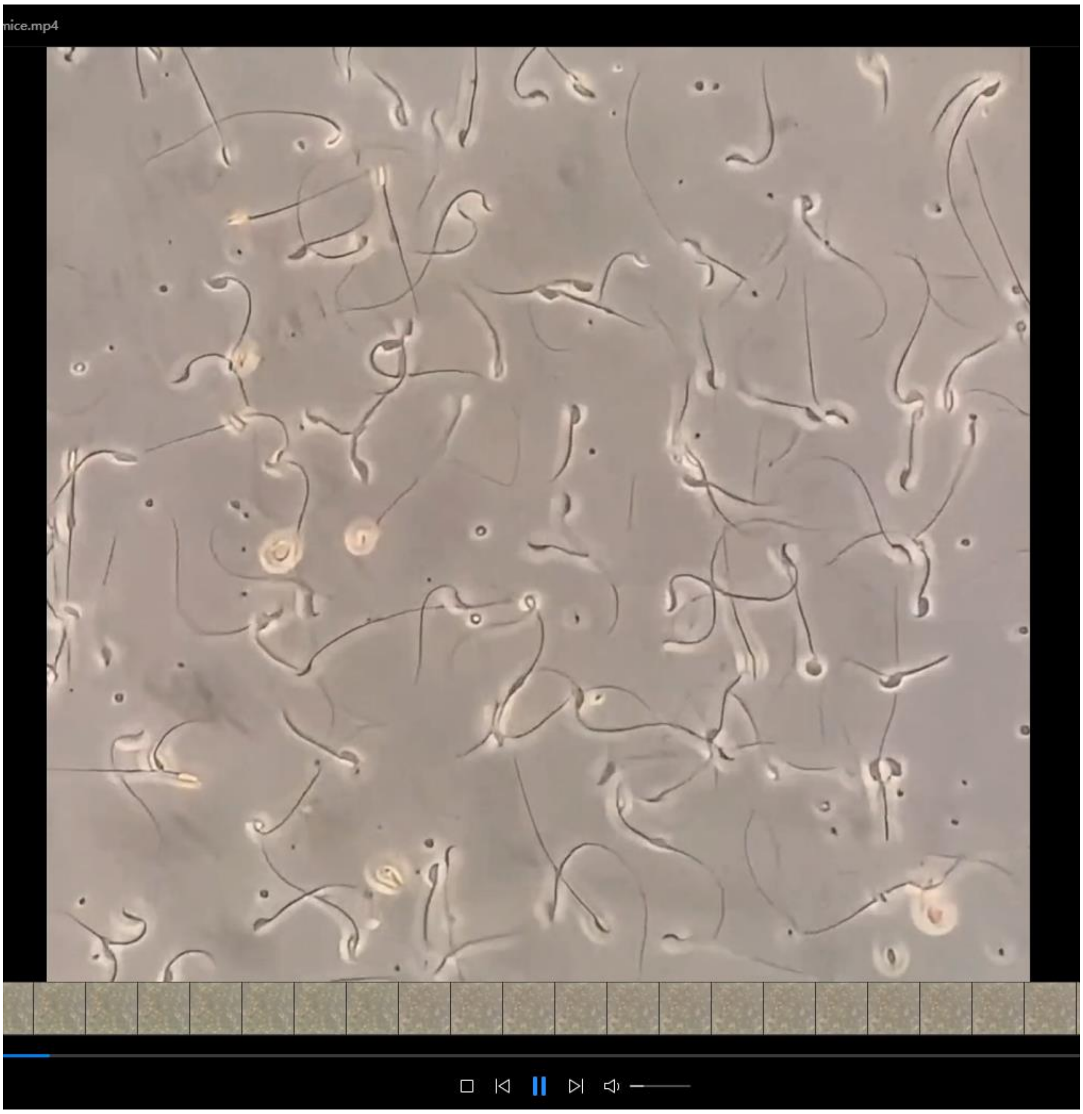


**Movie S1. Sperm motility of WT mice.**

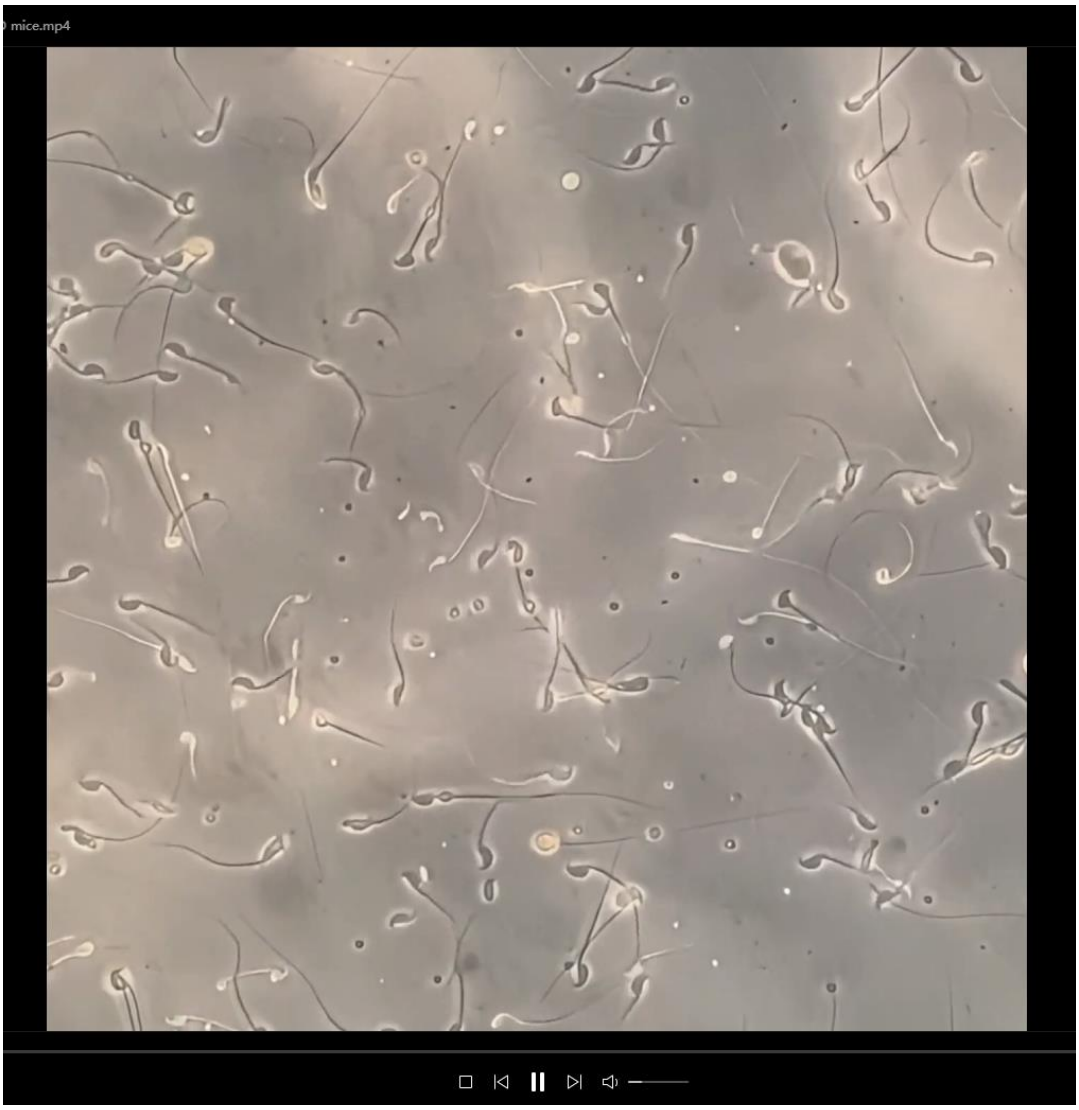


**Movie S2. Sperm motility of *Cfap77*-KO mice.**
